# Population genomics of *Candida albicans* reveals genetic footprints that allow the prediction of antifungal resistance

**DOI:** 10.64898/2026.10.28.762049

**Authors:** Adarsh Jay, Camille Bédard, Alicia Pageau, Isabelle Gagnon-Arsenault, Carla Bautista, Julianne V. Kus, Lisa McTaggart, Tanis C. Dingle, Sandy Shokoples, Domenica G De Luca, Amrita Bharat, Philippe J. Dufresne, Anna Fijarczyk, Christian R Landry

**Affiliations:** Département de Biochimie, de Microbiologie et de Bio-informatique, Faculté des Sciences et de Génie, Université Laval, Québec, QC G1V 0A6, Canada; Institut de Biologie Intégrative et des Systèmes, Université Laval, Québec, QC G1V 0A6, Canada; PROTEO, Le regroupement québécois de recherche sur la fonction, l’ingénierie et les applications des protéines, Université Laval, Québec, QC G1V 0A6, Canada; Centre de Recherche sur les Données Massives (CRDM), Université Laval, Québec, QC G1V 0A6, Canada; Département de Biologie, Faculté des Sciences et de Génie, Université Laval, Québec, QC G1V 0A6, Canada; Centre de Recherche en Infectiologie (CRI), Université Laval, G1V 0A6, Canada; Institut Intelligence et Données (IID), Université Laval, Québec, QC G1V 0A6, Canada; Public Health Ontario, Toronto, ON M5G 1M1, Canada; Department of Laboratory Medicine and Pathobiology, University of Toronto, Toronto, ON M5S 1A8, Canada; Alberta Precision Laboratories - Public Health Laboratory, Edmonton, AB T6G 2J2, Canada; National Microbiology Laboratory, Public Health Agency of Canada, Winnipeg, MB R3E 3R2, Canada; Department of Medical Microbiology and Infectious Diseases, University of Manitoba, Winnipeg, MB R3E 0J9, Canada; Laboratoire de santé publique du Québec, Institut national de santé publique du Québec Sainte-Anne-de-Bellevue, QC H9X 3R5, Canada; Département des Sciences Naturelles, Université du Québec en Outaouais, Ripon, QC J0V 1V0, Canada

## Abstract

Rapid detection of drug resistance in fungi is becoming increasingly urgent. While multiple loci and mutations were linked to antifungal resistance in *Candida albicans*, the utility of whole genome sequencing in predicting resistance has not been tested thoroughly. We address this by integrating more than 1000 *C. albicans* genomes, with associated resistance phenotypes to six antifungals, a curated repository of resistance-causing mutations and variant effect predictors. We used Genome Wide Association Studies and Machine Learning approaches to associate genetic variants to resistance phenotypes. We found that well curated databases like FungAMR are able to accurately identify markers predictive antifungal resistance. Curated mutations were detected in 56–74% of isolates with reduced azole susceptibility and 57–90% of those with reduced echinocandin susceptibility, revealing incomplete coverage of resistance across both drug classes. Complementary variant- and gene-level association analyses recovered established resistance determinants and uncovered additional candidate loci, whereas copy-number changes provided limited independent explanatory power. Genomic prediction declined when genetic backgrounds were absent from training, and when leading predictive features differed among backgrounds. We also found that models trained on a curated list of mutations and variant effect predictors performed well when classifying drug-susceptible strains, but the drug-resistance trait was predicted with a lower accuracy. These findings support the potential of genome-based resistance prediction while highlighting the need for broader population sampling, functional validation of candidate variants and expanded resistance catalogues to improve prediction across genetic backgrounds.

## Introduction

Invasive fungal diseases constitute a major and growing global-health challenge. The World Health Organization fungal priority pathogen list (WHO FPPL) was created in 2022 to guide research, development and public-health action against fungal infections and antifungal resistance. Laboratory capacity and surveillance are listed as core priorities^1^. *Candida albicans* is central to this problem because it is the most common agent of invasive fungal infections and a critical-priority pathogen in the WHO FPPL^1–3^.

Among the tools that would facilitate the surveillance of antifungal resistances is detection of resistance from whole-genome sequencing data. Comprehensive genome-based approaches could provide a fast alternative or complementary route to standard laboratory testing by identifying known resistance markers directly from sequence data. The utility of such markers in fungi still depends on the completeness of resistance variant catalogues and standardized bioinformatic interpretation, which are still limited^4,5^. Another limitation to a genomic approach is that *C. albicans* is highly diverse: the phylum contains multiple major and minor clades, with geographic or phenotypic overrepresentation in some clades, confounding drug resistance^6^. A third complication is that the *C. albicans* diploid genome itself is highly polymorphic but largely clonal^7^. Previous studies have shown that loss of heterozygosity and aneuploidy also contribute to its polymorphism^8,9^. In addition, copy-number variation has been shown to alter gene dosage and has been linked to antifungal resistance in the laboratory^10^, but its contribution to resistance across clinical fungal populations remains less clear compared to mutations^11^. This creates an unresolved problem for genomic surveillance: variants associated with the resistance phenotype may reflect shared ancestry rather than a functional contribution to resistance, while mutation catalogues may overlook contributions from structural variation.

Treatment options for systemic fungal infections of *C. albicans* remain limited, with only four main drug classes: polyenes, azoles, echinocandins and 5-flucytosine^12^. In *C. albicans*, azole resistance can arise through changes in the drug target Erg11, altered ergosterol biosynthesis, increased drug efflux or transcriptional rewiring^13^ while echinocandin resistance is more narrowly associated with substitutions in Fks1, the drug target of echinocandins^14^. More genes and mutations have been shown to be involved and curated resistance databases may provide sufficient data to interpret whole genome sequences. FungAMR is a manually curated catalogue containing 35,792 fungal mutations–drug records from 501 studies, 208 drugs, 246 genes and 95 species^15^. One can use its highest-confidence records (mutations tested directly in the native species or by expression in another organism, quantified through bulk competition assays, or supported by repeated emergence during experimental evolution), to ask how much resistance in a given genome could be assigned to established mechanisms. For *C. albicans* alone, this catalogue includes 6,604 records reporting resistance-associated changes for azoles and 179 for echinocandins. This provides an opportunity to determine which catalogued mutations occur in clinical populations and whether they are broadly represented or a smaller subset predominates among resistant isolates. It can also expose a likely gap: resistance in natural populations may be driven by rare variants, uncurated substitutions, combinations of mutations, copy-number changes or lineage-specific backgrounds.

Using a collection of whole-genome sequences from more than 1000 *C. albicans* isolates, we addressed four linked questions. First, we asked how completely current genome sampling captures the known population structure of *C. albicans*, and how large-scale genome changes are distributed across these populations, focusing on loss of heterozygosity (LOH), copy-number variation (CNV), aneuploidy and ploidy variation. Second, we asked how much antifungal resistance can be explained by fungAMR, the most comprehensive curated database available. Third, we asked whether genome-wide association analyses of single variants or CNVs can nominate additional candidate mechanisms. Finally, we asked whether whole-genome sequences can predict azole resistance, independent from phylogenetic associations.

## Results

### Population structure contextualizes genomic diversity and antifungal susceptibility

A total of 1173 *C. albicans* whole genome sequence dataset was assembled, and 1153 high quality genomes were retained for all downstream analysis (Methods, Genome processing; Strain genome quality control and Variant filtration). About half of the isolates come from seven published studies^8,16,18–22^ (n=626) and the other half were newly sequenced (n=527) (Fig 1a). The complete dataset includes strains isolated in 11 countries with Canada (n=685), China (n=370) and USA (n=54) contributing the most, followed by strains from Europe, other Asian countries and South Africa. We compiled MIC measurements reported in studies or generated by participating public health laboratories for six antifungals and classified them as binary resistant or susceptible phenotypes or recorded MICs when available. Throughout this study, isolates classified as susceptible were designated as the S phenotype, whereas those classified as intermediate, susceptible-dose-dependent or resistant were grouped as the R phenotype, representing reduced susceptibility (Methods, Antifungal susceptibility classification and multidrug-resistance definitions). The proportions of strains with resistance phenotypes in each group are summarized in Supplementary Table 1.

**Fig. 1.**
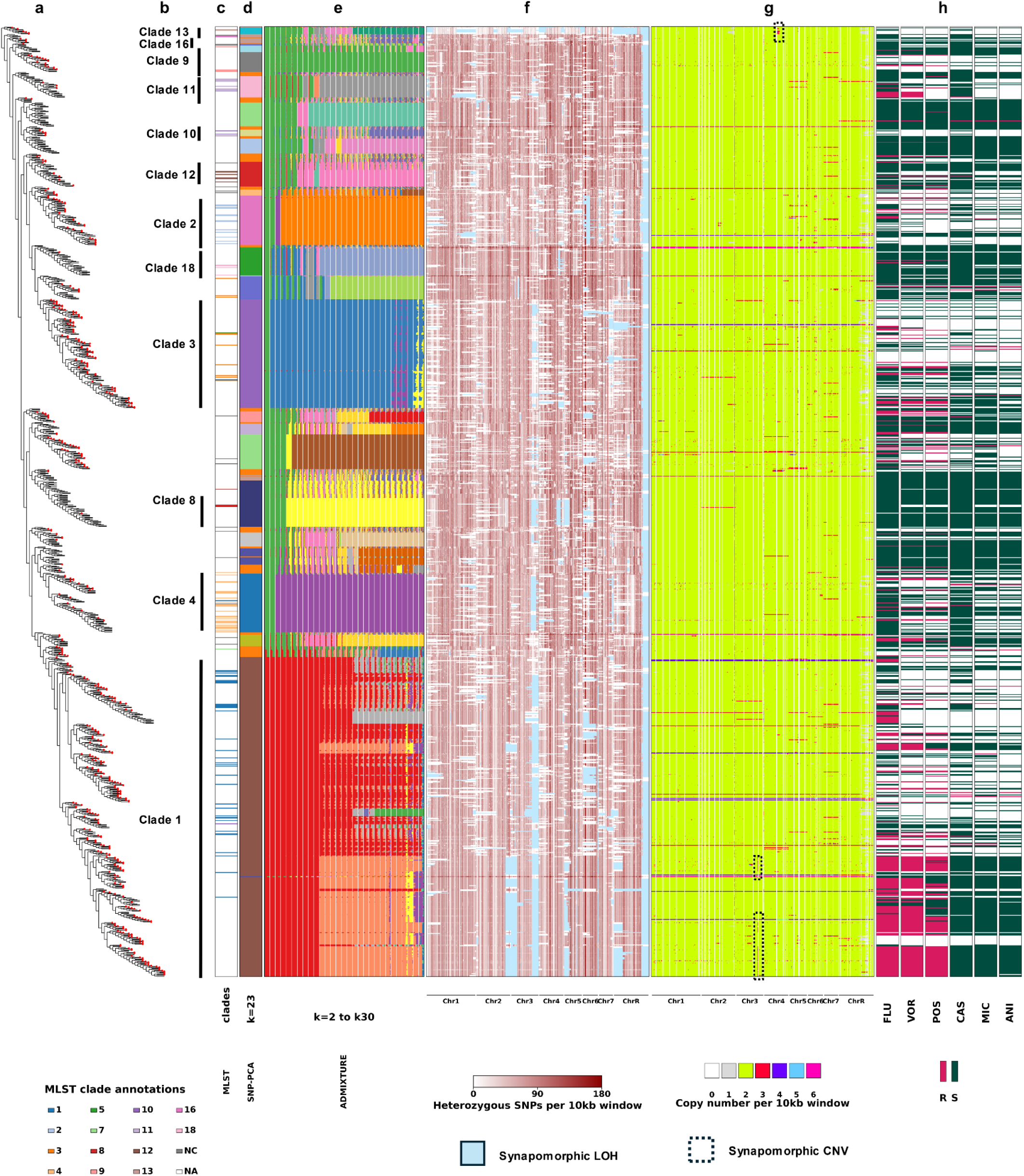
Integrated phylogenomic, admixture, loss-of-heterozygosity, copy-number and antifungal-resistance landscape across 1153 isolates of *C. albicans*. Each row represents one isolate, ordered identically across panels according to the phylogeny. **a,** Unscaled phylogenetic tree rooted using *Candida dubliniensis*. Newly sequenced isolates (n = 527) are marked in red. **b,** Clade labels derived from Multilocus sequence typing (MLST) (Methods, Clade annotation). **c,** MLST assignments used as references for clade annotation^16,17^. NC denotes the new clade described by Sitterle et al.^16^; NA denotes isolates without a published MLST annotation. **d,** Membership of the 23 clusters identified by k-means clustering of principal-component coordinates. **e,** ADMIXTURE ancestry profiles for K = 2–30, displayed as 29 adjacent columns, with one column per K. The replicate with the lowest cross-validation error was selected for each K. Colours identify ancestry, and stacked segments indicate their proportions in each isolate. **f,** Loss of heterozygosity (LOH), represented by the number of heterozygous single-nucleotide polymorphisms per 10-kb window. Darker red indicates greater heterozygosity. Light-blue marks phylogenetically shared LOH segments (Methods, LOH detection and quantification). **g,** Genome-wide copy number per 10-kb window. Colours indicate copy-number states from 0 to 6; dashed outlines identify phylogenetically shared copy-number changes. Columns in the heatmaps in **f** and **g** follow chromosome order along the *C. albicans* SC5314 haplotype A reference genome (Chr1–Chr7 and ChrR) **h,** Experimental binary susceptibility phenotypes for fluconazole (FLU), voriconazole (VOR), posaconazole (POS), caspofungin (CAS), micafungin (MIC) and anidulafungin (ANI) (Methods, Antifungal susceptibility classification and multidrug-resistance definitions). Magenta indicates isolates classified as R and green indicates isolates classified as S under the study-defined thresholds; white indicates that no data is available on R and S. Keys below the panels define clade and cluster colours, heterozygosity and copy-number scales, and shared-segment annotations.

A maximum-likelihood phylogeny resolved multiple deeply branching lineages (Methods, Phylogenetic Tree Building; Fig. 1a). Previous multilocus sequence typing (MLST) studies established a framework of 18 numbered clades of *C. albicans^7^*, with an additional clade subsequently reported by Sitterlé et al. (2020)^16^, referred to as NC. These clades represent groups of genetically related isolates, and published MLST annotations provided labels for 139 isolates in our collection, spanning 14 numbered clades and NC (Fig. 1c). To characterize genetic structure across the full collection, including isolates without published clade labels, we performed principal-component analysis (PCA) followed by unsupervised clustering. This identified 23 genetic groups that generally matched the phylogenetic divisions and were robust to resampling (Methods, Linkage disequilibrium (LD) pruning of the SNP dataset; Principal Component Analysis (PCA) of SNPs; Unsupervised clustering; Fig. 1a,d; Extended Data Fig. 1).

These groups agreed much more closely with published MLST clades than with geographic origin, contributing study or fluconazole resistance, showing that the genetic divisions did not simply reproduce the sampling or resistance categories (Extended Data Fig. 2).

We next used ADMIXTURE to estimate the ancestry composition of each isolate, varying the number of ancestry components allowed in the model, denoted K, from 2 to 30. The ancestry profiles supported the major clades and revealed finer subdivisions within them as K increased (Methods, ADMIXTURE analysis; Fig. 1e). The model’s ability to predict withheld genetic data improved relatively little beyond K = 23, although prediction error was lowest at K = 29 (Extended Data Fig. 1g). Published MLST labels showed strong overlap with PCA clustering: 128 of 139 isolates (92.1%) matched the MLST label within their PCA cluster (Extended Data Fig. 2a). Phylogenetic placement and ancestry profiles were also broadly consistent with these assignments (Extended Data Fig. 2b). This agreement supported extending established clade names to unlabelled isolates using PCA-cluster boundaries, while retaining cluster labels where a named-clade assignment was unsupported (Methods, Clade annotation; Fig. 1b,d; Extended Data Fig. 2a,b). Most assigned clades remained recognizable across ancestry resolutions, whereas clade 1 separated into several sublineages (Fig. 1b,e). We therefore refer to genetic backgrounds by their assigned clade name or, where unavailable, their PCA-cluster number. Overall, the sequencing of more than 500 additional genomes resolved more than 20 different genetic backgrounds that covers much of the known genetic diversity of *C. albicans*, while also revealing population structure beyond that captured by established clades.

Large-scale genome variation also differed among genetic backgrounds. Loss of heterozygosity (LOH) was detected in 1,152 isolates (99.9%). To assess whether local deletions could explain these patterns, we compared LOH tracts with regions of copy-number loss. These rarely overlapped, suggesting that local deletions accounted for only a small proportion of the observed LOH (Methods, LOH detection and quantification; Fig. 1f,g). The extent of LOH varied substantially among clades, with clade 13, also known as *Candida africana*, showing the greatest incidence of LOH and clade 9 the least. LOH tracts frequently extended towards chromosome ends. Isolates within the same clade often shared LOH tracts at similar genomic positions, suggesting that some tracts were inherited from a common ancestor (Fig. 1f; Supplementary Table 2). The frequent sharing of LOH tracks among strains suggest that they are not only present because of their high rate of occurrence but also because they are maintained in populations.

Copy-number variation (CNV) was defined as copy-number gains or losses in genomic regions ≥30 kb in length, relative to baseline ploidy. Genome coverage and allele frequency analyses revealed that most isolates were diploid (n=1127, 97.7%), but a small minority showed triploid (n=8, 0.7%), tetraploid (n=11, 1.0%) or putative hexaploid (n=7, 0.6%) profiles (Methods, Whole-genome ploidy and CNV detection; Extended Data Fig. 3a–d). This diversity of ploidy is consistent with other recent reports showing that ploidy in this species is highly variable also within a geographical region ^23^. LOH patterns of triploid and tetraploid isolates matched that of their phylogenetic neighbours while putative hexaploids had markedly less LOH than all other isolates (Fig. 1f). Whole-chromosome copy-number changes occurred most frequently on chromosome 7, were exclusively gains of chromosomes (n=55), and were all trisomies except for one tetrasomy in chromosome 7. Segmental changes (CNVs of regions smaller than a whole chromosome) included both gains and losses (Extended Data Fig. 3e,f). Shared segmental gains on chromosome 3 within clade 1 and chromosome 4 within clade 13 provided further examples of genome changes associated with particular lineages (Fig. 1b,g).

Resistance differed sharply between drug classes (Fig. 1h; Extended Data Fig. 4a). R phenotype frequencies differed sharply between drug classes (Fig. 1h; Extended Data Fig. 4a). The fluconazole R phenotype occurred in 61.6% of tested clade 1 isolates (167/271), compared with 16.2% outside clade 1 (97/600). Clade 1 accounted for 63.3% of all fluconazole R phenotype isolates (167/264), although this phenotype also occurred in seven other named clades and additional genetic backgrounds (Fig. 1a,b,h). Echinocandin R phenotypes were uncommon and distributed across several genetic backgrounds (Fig. 1b,h). Across the collection, R phenotype frequencies among tested isolates were 30.3% for fluconazole (264/871), 33.4% for voriconazole (220/659) and 28.6% for posaconazole (175/611), compared with 3.0% for caspofungin (23/766), 2.4% for micafungin (16/667) and 2.0% for anidulafungin (12/589; Extended Data Fig. 4a). Within-class cross-resistance, defined as an R phenotype for at least two drugs, occurred in 34.0% of isolates tested against at least two azoles (224/659), compared with 2.4% of those tested against at least two echinocandins (15/632). Among isolates tested against all three drugs within each class, 26.6% had R phenotypes for all three azoles (162/610), compared with 1.9% for all three echinocandins (11/575). Multidrug resistance spanning both classes occurred in 1.2% of isolates tested against at least one drug from each class (10/809; Extended Data Fig. 4b).

Together, these observations reveal a strongly structured population with uneven distributions of resistance and genome variation, implying a need to assess resistance-associated variants across genetic backgrounds.

### Curated resistance mutations account for the majority of the observed R phenotype

We next assessed how well curated known resistance mutations could explain the phenotypes in our *C. albicans* collection. We screened *ERG11, UPC2, CDR1, CDR2, MDR1, TAC1, MRR1, MRR2, CAP1* for azole resistance and *FKS1* for echinocandin resistance, encompassing drug targets, efflux pumps and resistance regulators^14,26–29^. Across these genes, the curated database FungAMR contained 1218 amino acid substitutions in 5 genes (*ERG11, UPC2, MRR1, TAC1* and *FKS1)* associated with resistance to the six antifungal drugs in this study with relatively high-confidence (Methods, FungAMR annotation and antifungal susceptibility). Most observed amino-acid substitutions lacked matching FungAMR records, and database matches were concentrated in Erg11, indicating uneven coverage of variation across these established resistance-associated genes (Extended Data Fig. 5a; Supplementary Table 3).

Common high-confidence Erg11 resistance substitutions showed pronounced phylogenetic clustering. A114S and Y257H always co-occurred and were restricted to clade 1, whereas T123I occurred exclusively with Y132H, predominantly within a distinct clade 1 sublineage that becomes visible at ADMIXTURE ancestry proportions k=28 (Methods, ADMIXTURE analysis; Fig. 2a; Extended Data Fig. 5d). These patterns previously reported in Chinese isolates^20^, were also present in our newly sequenced Canadian isolates. High-confidence substitutions in resistance regulators (Upc2 A643V and G648D, Tac1 N977D and G980E, and Mrr1 Q350L and P683S) were less frequent. Fks1 substitutions were uncommon: S645P and R647G occurred across several genetic backgrounds, whereas F641S was confined to clade 1 (Fig. 2a; Extended Data Fig. 5d). Together, these patterns show recurrent, clade-associated combinations of common Erg11 substitutions, with rarer Fks1 substitutions occurring across multiple genetic backgrounds.

**Fig. 2.**
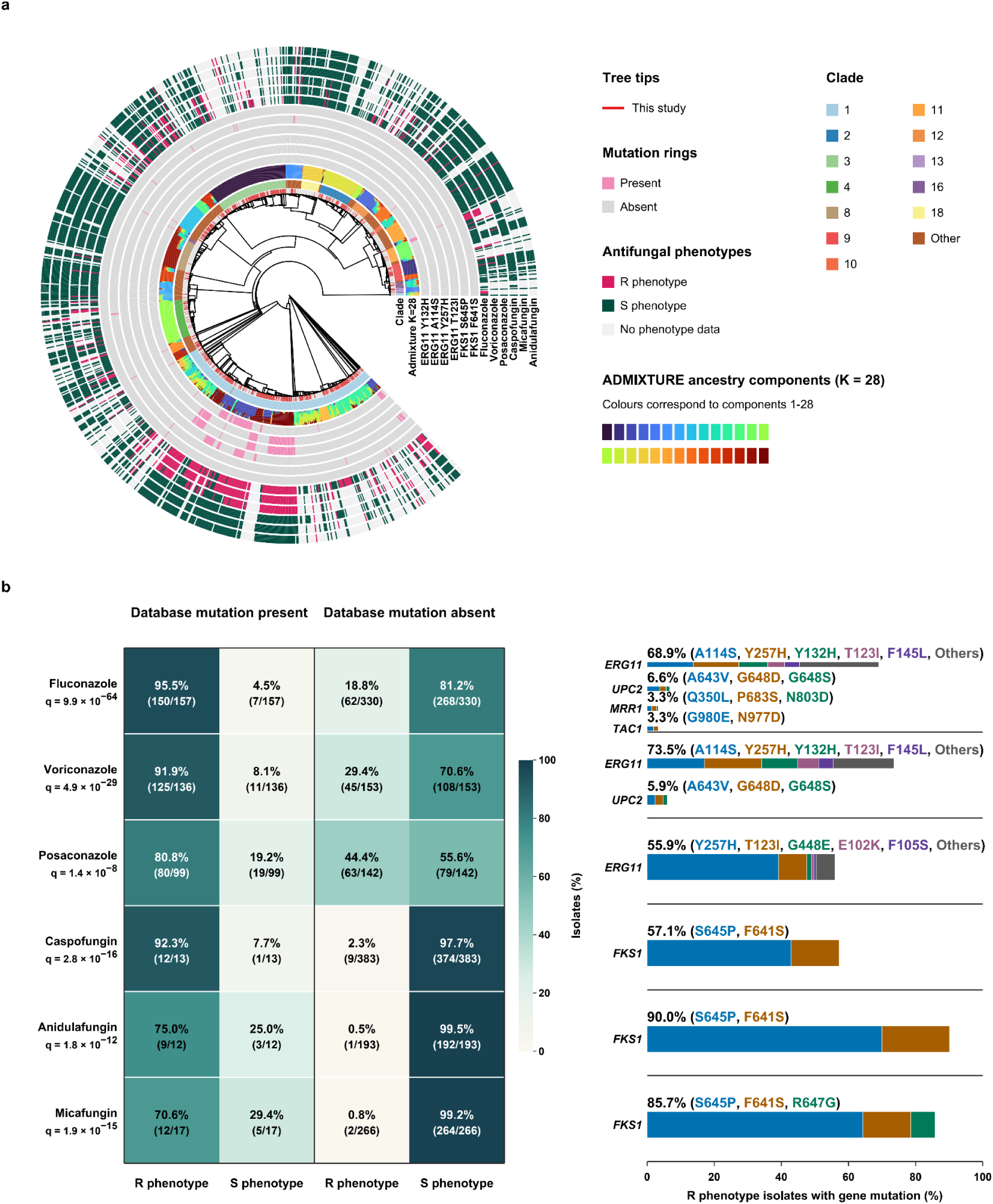
Phylogenetic distribution and phenotypic concordance of high-confidence resistance associated with amino acid substitutions. **a,** Maximum-likelihood phylogeny of 1,153 *C. albicans* isolates. Red tips mark isolates sequenced in this study. Rings show clade, ADMIXTURE ancestry proportions (*K* = 28), Erg11 and Fks1 substitutions, and antifungal R and S phenotypes. Pink and grey indicate substitution presence and absence; magenta, dark green and light grey indicate R phenotype, S phenotype and unphenotyped isolates, respectively (Methods, Antifungal susceptibility classification and multidrug-resistance definitions). Branch lengths are displayed schematically. **b,** Association between drug-matched high-confidence substitutions (resolved FungAMR confidence scores +1 to +4) and R/S phenotype, using allele-specific annotations and alternate-allele carriage from the standardized variant catalogue. Marker definitions and signed-evidence resolution are described in (Methods, FungAMR annotation and antifungal susceptibility). Isolates from Gong et al. (2023)^20^ (n = 370) and Chew et al. (2021)^18^ (n = 14) were excluded from this assessment because these studies contributed data to FungAMR. The remaining assessment includes 494 isolates with at least one binary phenotype: n = 487, 289, 241, 396, 205 and 283 for fluconazole, voriconazole, posaconazole, caspofungin, anidulafungin and micafungin, respectively. Heatmap percentages are calculated separately within database-mutation-present and database-mutation-absent groups for each drug; counts and group denominators are shown in parentheses. Database mutation absence means no detected qualifying drug-matched substitution. Each gene bar shows the percentage of all isolates with R phenotype for that drug carrying at least one qualifying substitution in that gene; carrier sets for different genes can overlap. Within a gene, each isolate contributes an equal weight of 1/k to each of its k qualifying substitutions. The five substitutions with the largest weighted contributions are shown individually, with the remainder grouped as Others. Colours follow the same substitution-rank sequence in every gene bar, with Others in grey. Coloured substitution names match their stacked-bar segments; these weights partition the gene total. Associations were tested using two-sided Fisher’s exact test^24^ on database mutation presence/absence × R phenotype/S phenotype contingency tables. Values labelled q are P values adjusted across all six drugs using the Benjamini–Hochberg procedure (1995)^25^. Exact P values, odds ratios, contingency tables and substitution weights are provided in the source data.

Allelic states also differed among resistance mutations. Common Erg11 and Fks1 substitutions were predominantly homozygous, and their genes frequently overlapped LOH tracts. *MRR1* overlapped a called LOH tract in all carriers of Q350L and P683S, whereas Upc2 A643V and G648D were exclusively heterozygous and showed no gene-overlapping LOH. (Methods, FungAMR annotation and antifungal susceptibility; LOH detection and quantification; Extended Data Fig. 5b,c; Supplementary Table 3). Thus, homozygosity and LOH frequently accompanied substitutions previously characterized to contribute to resistance, but not always.

Despite the lack of complete coverage, the database likely covers the most frequently encountered resistance mutations. We tested whether isolates carrying high-confidence FungAMR mutations were more likely to be resistant to the drugs to which these mutations have been associated with (Methods, FungAMR annotation and antifungal susceptibility). Across 494 isolates with available phenotypes, substitutions were significantly associated with the R phenotype for all six antifungals (two-sided Fisher’s exact tests with Benjamini–Hochberg adjustment; all q ≤ 1.4 × 10⁻⁸; Fig. 2b; Supplementary Table 3). Among substitution carriers, the proportion with the R phenotype was 95.5% for fluconazole (150/157), 91.9% for voriconazole (125/136), 80.8% for posaconazole (80/99), 92.3% for caspofungin (12/13), 75.0% for anidulafungin (9/12) and 70.6% for micafungin (12/17). Erg11 A114S and Y257H predominated among fluconazole- and voriconazole-associated substitutions, with Y132H also prominent, whereas Y257H predominated for posaconazole. The leading substitutions in regulatory genes were Upc2 A643V for fluconazole and A643V and G648D for voriconazole, alongside Mrr1 Q350L and P683S and Tac1 G980E and N977D for fluconazole. Echinocandin-associated substitutions were concentrated in Fks1, principally S645P and F641S. Among isolates without a qualifying drug-matched substitution, the R phenotype was observed in 18.8%, 29.4% and 44.4% of isolates tested against fluconazole, voriconazole and posaconazole, respectively, compared with 2.3%, 0.5% and 0.8% for caspofungin, anidulafungin and micafungin (Fig. 2b). The complete list of FungAMR-matched substitutions detected across our genome collection, is provided in Supplementary Table 3. Thus, curated mutations were strong indicators of resistance (>=90%), but substantial gaps still exist in our ability to identify resistant isolates using currently catalogued mutations. Many more unreported mutations likely contribute to resistance to azoles while Fks1 substitutions seem to explain the majority of echinocandin resistance.

Resistance has been associated with CNVs, particularly often in laboratory experiments^11^. Therefore, we examined whether the presence of CNVs could help explain resistance across our collection. Among diploid isolates (n=1127), CNVs most frequently affected *ERG11* and *TAC1* and were almost exclusively copy-number gains (Methods, Whole-genome ploidy and CNV detection; Extended Data Fig. 6a,b; Supplementary Table 2). These included chromosome 5L amplifications encompassing both genes, with profiles consistent with the azole-associated i(5L) structure^10^ (Extended Data Fig. 6c). However, after excluding high-confidence mutation carriers, no *ERG11* or *TAC1* gain carrier was resistant to the tested azoles among those phenotyped (Methods, Copy-number variation and antifungal susceptibility; Extended Data Fig. 6d; Supplementary Table 2). *UPC2* gains accompanied the largest increases in median voriconazole and posaconazole MIC, but each comparison involved only two eligible carriers (Methods, Copy-number variation and antifungal susceptibility; Extended Data Fig. 6e; Supplementary Table 2). Evidence that these CNVs contribute to resistance by themselves is therefore limited, as recently shown in our metaanalysis^11^.

The relationship between *ERG11* amplification and MIC varied across backgrounds defined by mutations in *ERG11*. Gains accompanied higher azole MICs in A114S/Y257H and G448E carriers for particular drugs, whereas Y132H carriers showed no change and Y132F carriers showed lower fluconazole MICs (Methods, Copy-number variation and antifungal susceptibility; Extended Data Fig. 6f; Supplementary Table 2). The small, partly overlapping groups of two to four CNV-plus-mutation carriers support a possible modifying role for gene dosage, without establishing synergy.

Together, these results support curated substitutions as useful but incomplete markers of resistance. Qualifying drug-matched substitutions were absent from 29.2% of fluconazole, 26.5% of voriconazole and 44.1% of posaconazole R-phenotype isolates, indicating substantial variation in marker coverage among drugs. Copy-number gains were associated with differing MICs in some ERG11 substitution backgrounds, but the small, partly overlapping carrier groups limit inference about their independent contribution to resistance.

### Genome-wide association studies reveal established and candidate genes associated with azole and echinocandin resistance

We tested individual SNPs and indels for associations with antifungal susceptibility, measured by log₂MIC, to identify candidate resistance mechanisms beyond those catalogued in FungAMR (Methods, Genome-wide Association Study of SNPs and Indels; Fig. 3a; Supplementary Tables 4–9). The strongest signals recovered the established drug targets *ERG11* and *FKS1^14,30^*. Erg11 K143R (P = 9.70 × 10⁻¹¹), Y132H (P = 2.19 × 10⁻⁸) and F126L (P = 2.68 × 10⁻⁸) were significant for fluconazole, with K143R showing the strongest association. Erg11 G450E (P = 1.70 × 10⁻⁷) and R467K (P = 6.05 × 10⁻⁷) and Tac1 E461K (P = 8.08 × 10⁻⁷) met a less stringent exploratory threshold (Supplementary Table 4). Erg11 Y132H was the strongest voriconazole association, whereas no *ERG11* variant reached genome-wide significance for posaconazole (Supplementary Tables 5,6). Fks1 S645P (caspofungin, P = 3.85 × 10⁻³²; micafungin, P = 7.87 × 10⁻⁵³; anidulafungin, P = 1.25 × 10⁻²⁴) was the strongest association for all three echinocandins, indicating a more consistent target-gene signal across this drug class (Supplementary Tables 7–9).

**Fig. 3.**
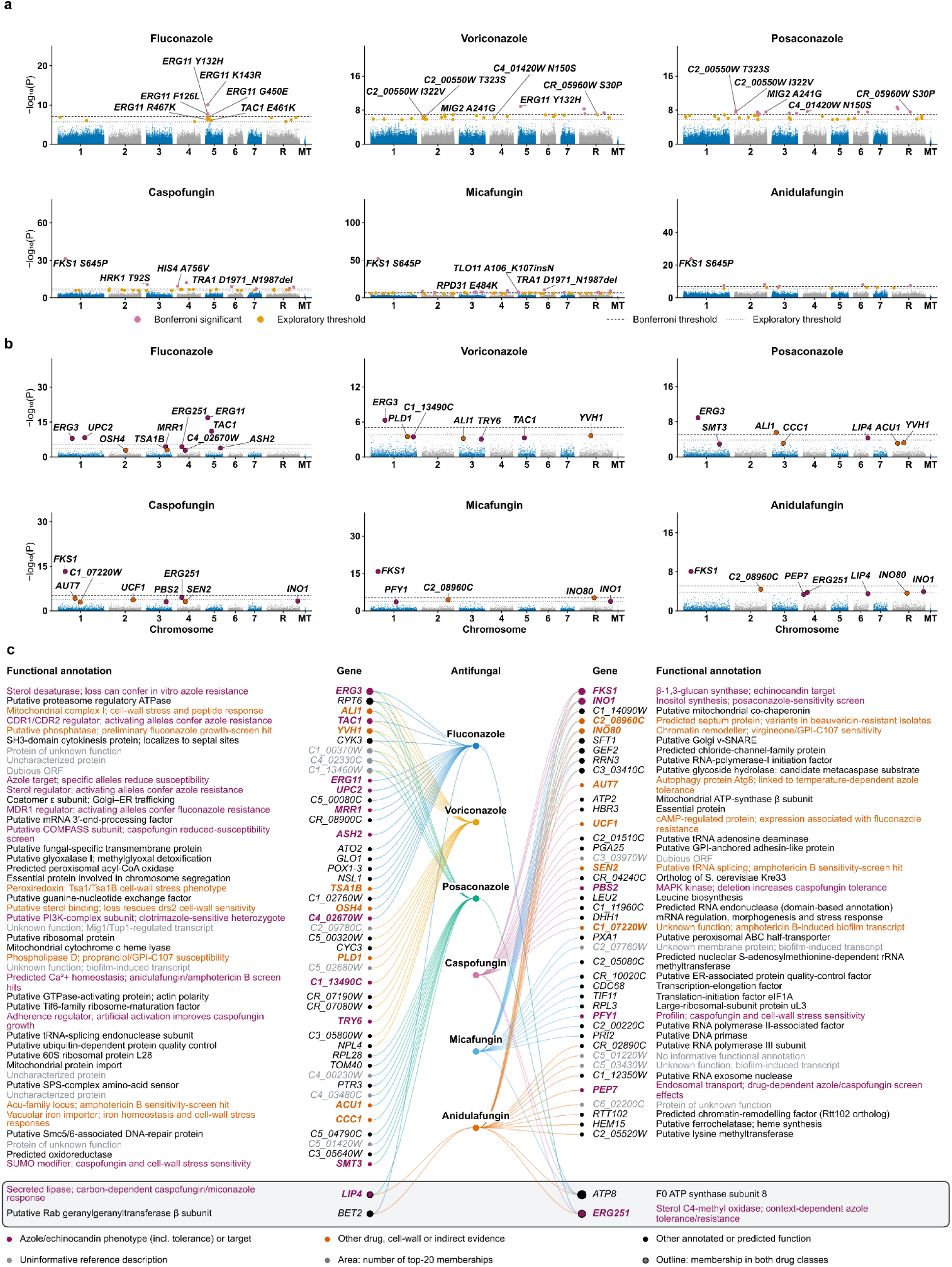
Variant- and gene-level associations with antifungal susceptibility. **a,** Associations of individual SNP and indel alleles with log₂MIC. Each point represents one allele. Pink and orange indicate Bonferroni-significant and suggestive associations, respectively; selected variants are labelled. **b,** Associations of gene-level carrier indicators formed from variants with predicted HIGH or MODERATE impact carried by ≤5% of each drug-specific cohort. Each point represents one tested gene. Enlarged magenta and orange points identify literature-supported genes among the 20 highest-ranked associations for each drug; these genes are labelled. Analyses included 871 isolates for fluconazole, 659 for voriconazole, 611 for posaconazole, 766 for caspofungin, 667 for micafungin and 589 for anidulafungin. In **a** and **b**, alternating blue and grey distinguish chromosomes, including the mitochondrial genome (MT). Dashed and dotted lines indicate the drug-specific Bonferroni (*P* = 0.05/*M*) and suggestive (*P* = 1/*M*) thresholds, where *M* is the number of tested alleles or genes. *P* values were obtained from two-sided mixed-model association tests. **c,** Network of the 20 highest-ranked genes per drug, comprising 91 distinct genes and 120 drug–gene memberships. Edge colours identify drugs. Gene-marker area scales with the number of top-20 memberships; outlined markers identify genes represented in both antifungal classes. In **b** and **c**, magenta denotes published azole/echinocandin phenotype or drug target evidence, and orange denotes evidence involving other novel agents, cell-wall phenotypes or indirect drug responses. In **c**, black denotes other annotated or predicted functions, and grey denotes uninformative functional annotations.

Associations outside these targets suggest the implication of additional cellular processes. Posaconazole-associated variants occurred in *MIG2*, which contributes with *MIG1* to carbon metabolism and cell-wall stress responses^31^. Associations shared between posaconazole and voriconazole also occurred in the poorly characterized loci *C2_00550W_A*, *C4_01420W_A* and *CR_05960W_A*; these reached genome-wide significance for posaconazole and the exploratory threshold for voriconazole (Fig. 3a; Supplementary Tables 5,6). Echinocandin associations included the transcriptional regulators *TRA1*, *RPD31* and *TLO11*, which have published cell-wall or antifungal-response evidence^32–34^, alongside *HRK1* and *HIS4*, involved in ion balance and histidine synthesis, respectively^35,36^ (Fig. 3a; Supplementary Tables 7,8).

De novo mutations causing resistance are expected to be rare in populations. Because rare variants can be difficult to detect and can hardly show strong association with phenotypes, we combined variants with predicted HIGH or MODERATE functional impact carried by ≤5% of each drug-specific cohort into gene-level carrier indicators^37^ (Methods, Gene-level variant GWAS; Fig. 3b; Supplementary Tables 10–15). This recovered additional established determinants: *ERG11*, *TAC1*, *UPC2*, *ERG3* and *C5_00080C* were associated with higher fluconazole MICs at genome-wide significance (Supplementary Table 10). Aggregation therefore strengthened the evidence for *TAC1* and recovered the established azole-resistance genes *UPC2* and *ERG3^38–40^*. *ERG3* was significant across all three azoles and was the only significant gene for voriconazole (Supplementary Tables 10–12). *FKS1* remained significant across all three echinocandins and was the only significant gene for anidulafungin (Supplementary Tables 13–15). Additional significant candidates included *ALI1* for posaconazole that encodes a mitochondrial NADH-ubiquinone oxidoreductase subunit, *C1_14090W* for caspofungin and micafungin, and *INO80* for micafungin a core catalytic subunit involved in chromatin-remodeling (Supplementary Tables 12–14); published screening evidence links *INO80* to susceptibility to the experimental antifungal compound GPI-C107^41^. Quantile–quantile plots for the variant- and gene-level scans are shown in Extended Data Fig. 7a,b.

At the exploratory threshold (P<1.61 × 10⁻⁴), the efflux regulator *MRR1* substitutions were associated with higher fluconazole MIC^42^, while *RPT6* was associated with all three azoles (Fig. 3b; Supplementary Tables 10–12). Other associations involved *ASH2* and *INO1*, previously linked to antifungal susceptibility in screening studies^41^; *AUT7*, implicated in autophagy-dependent azole tolerance^43^; and *C2_08960C*, previously reported alongside *TAC1* mutations in beauvericin-resistant isolates^44^ (Fig. 3b; Supplementary Tables 10, 13–15). These findings broadened the candidate set to include regulatory and stress-response functions.

Four genes connected the top-20 lists across antifungal classes (Methods, Network of leading gene-level variant associations; Fig. 3c; Supplementary Tables 10, 12–15). *ERG251* appeared for fluconazole, caspofungin and anidulafungin, and *LIP4* for posaconazole and anidulafungin, providing links to sterol metabolism and lipid utilization^45,46^. The mitochondrial gene *ATP8* appeared for posaconazole and all three echinocandins, but its anidulafungin association at the exploratory threshold involved only one carrier (Supplementary Table 15). *BET2* appeared for fluconazole and anidulafungin without meeting the exploratory threshold for either (Supplementary Tables 10,15). These overlaps identify candidates for further investigation for cross-resistance.

Other highlighted genes did not meet the exploratory threshold but had relevant published evidence (Fig. 3b,c; Supplementary Tables 10–15). These included *PFY1* and *SMT3* for caspofungin sensitivity^47,48^, and *TSA1B* for cell-wall phenotypes following combined disruption with *TSA1^49^*. Deletion of *PBS2* increases caspofungin tolerance^50^. *SEN2* was identified in an amphotericin B sensitivity screen^51^, while *OSH4* deletion suppresses calcofluor-white sensitivity in a *drs2* mutant background^52^. Antifungal screening evidence supported *PEP7*, *C1_13490C*, *ACU1* and *PLD1^41^*, while *C4_02670W* and *CCC1* were linked to clotrimazole susceptibility and iron homeostasis, respectively^53,54^. Expression studies implicated *UCF1* in acquired azole resistance and *C1_07220W* in responses to amphotericin B^55,56^, while functional screens linked *YVH1* to fluconazole responses and *TRY6* to growth under caspofungin exposure^57,58^.

Together, individual-variant and gene-level analyses recovered established resistance genes as contributing strongly to resistance but also revealed many other genes potentially impacting drug susceptibility. While many of these genes have previously been associated with resistance, some show new associations. Understanding their role in antifungal resistance will require further study.

### Population structure explains most of the apparent machine-learning predictability of fluconazole resistance

Combinations of genetic variants may help predict resistance even when their individual associations are difficult to detect. We therefore used machine learning to test how well genomic variation predicted resistance across genetic backgrounds and to compare the variants and genes most useful for prediction with those identified by GWAS.

We compared random forests (RF) using three feature sets: individual variant dosages, recording the number of copies of each alternate allele; gene-level variant profiles, recording whether each gene carried a rare variant predicted to affect its function; and phylogeny-based features comprising SNP-derived principal components (PCs). The phylogeny-based approach therefore used summaries of genomic relatedness without receiving individual variants or genes as separate predictors. Analyses included 594 isolates from seven genetic backgrounds, each containing ≥15 isolates and ≥5 resistant isolates (Methods, Random-forest prediction of fluconazole resistance; Fig. 4a; Supplementary Table 16).

**Fig. 4.**
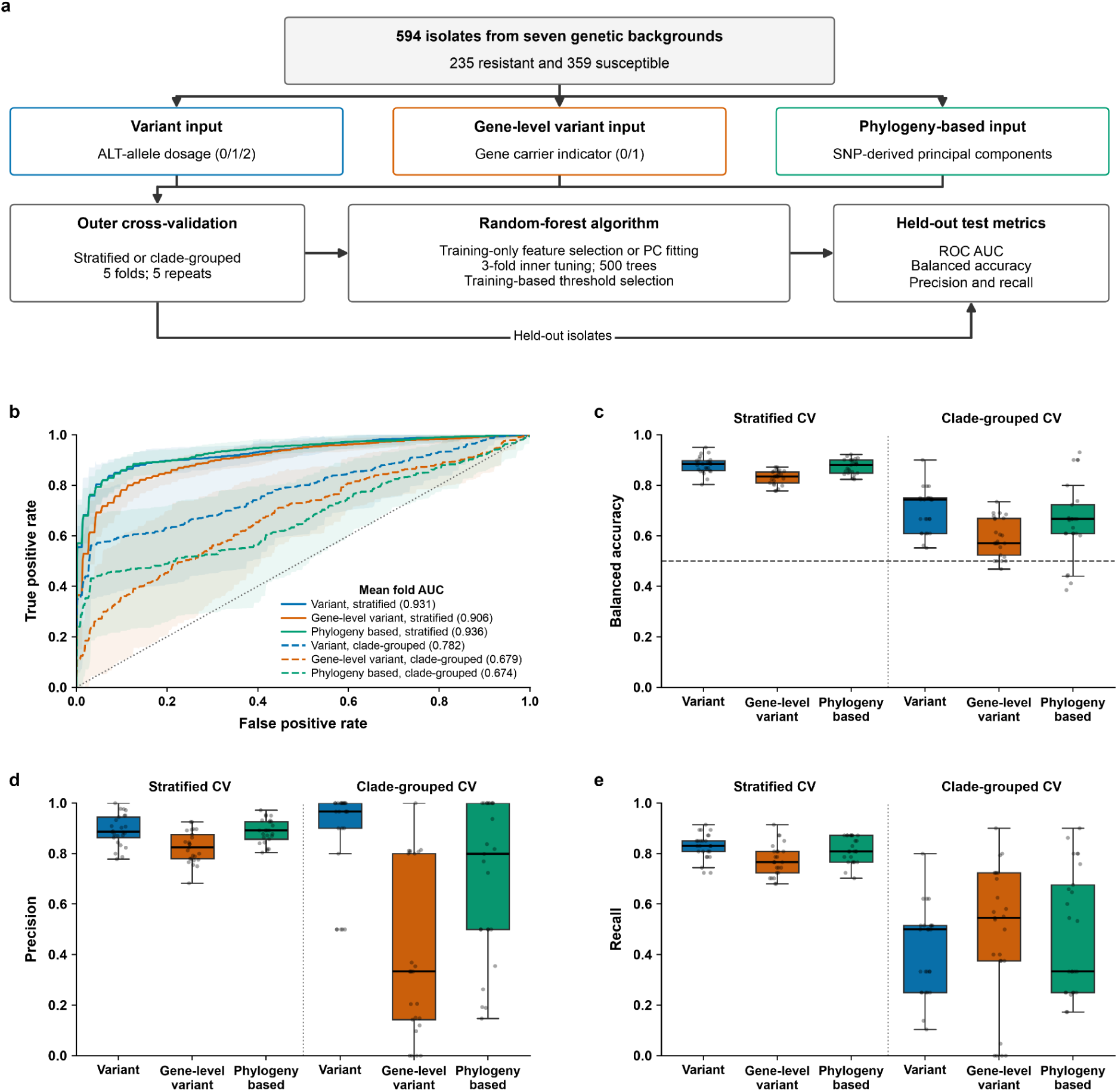
Random-forest prediction of fluconazole resistance across seven genetic backgrounds. **a,** Workflow for 594 isolates (235 resistant and 359 susceptible) from seven different genetic backgrounds containing ≥15 isolates and ≥5 resistant isolates. Inputs derive from one standardized SNP-and-indel catalogue and comprise alternate-allele dosages (0/1/2), binary gene-level indicators of variants with predicted HIGH or MODERATE impact and phylogeny-based principal components (PCs) derived from the common linkage disequilibrium-pruned nuclear SNP panel also used for GWAS relatedness estimation. Each input was evaluated using five-fold stratified or clade-grouped cross-validation (CV) with five repeats. Feature selection, PC fitting, tuning and threshold selection were restricted to training data. **b,** Mean receiver operating characteristic (ROC) curves for variant-based (blue), gene-level variant-based (orange) and phylogeny-based (green) predictions. Solid and dashed lines denote stratified and clade-grouped CV, respectively. Curves are averaged across 25 outer folds; shading indicates ±1 s.d. Key values give mean outer-fold area under the ROC curve (AUC). The dotted diagonal indicates no discrimination. **c–e,** Balanced accuracy (**c**), precision (**d**) and recall (**e**) across the same 25 outer folds. Colours identify inputs. Boxes span the first and third quartiles, centre lines mark medians and whiskers extend to the furthest observations within 1.5 times the interquartile range beyond either quartile. Points show all fold estimates, including outliers. Vertical dotted lines separate validation schemes; the horizontal dashed line in **c** marks balanced accuracy of 0.5. Balanced accuracy averages resistant- and susceptible-class recall. Precision and recall concern the resistant class; precision is zero when no resistant isolates are predicted. Grouped CV keeps each genetic background within training or testing. The same five held-out group combinations recur across repeats, with varying fitting and inner-validation seeds.

Under stratified cross-validation (CV), where genetic backgrounds were represented in both training and test sets, phylogeny-based and variant-based predictions performed similarly. Their mean receiver operating characteristic area under the curve (ROC AUC) was 0.936 and 0.931, respectively, compared with 0.906 for gene-level predictions (Fig. 4b; Supplementary Table 16). Variant-based predictions achieved slightly higher balanced accuracy, precision and resistant-isolate recall, while gene-level predictions performed less well across these measures (Fig. 4c–e). Thus, phylogeny-based predictions performed about as well as variant-based predictions, consistent with the observation of resistance mutations being more common in some clades than others, making genetic background itself informative for predicting the R phenotype.

To assess whether resistance predictions generalize to clades not represented in the training data, we repeated validation while keeping isolates from each clade entirely within either the training or test set (Extended Data Fig. 8a). Performance declined for all three feature sets, but variant-based predictions retained the highest mean ROC AUC (0.782), compared with gene-level (0.679) and phylogeny-based predictions (0.674; Fig. 4b; Supplementary Table 16). Variant-based predictions also achieved the highest balanced accuracy and precision. However, the models identified only a mean fraction of 0.416 of isolates with the R phenotype, despite a high mean precision of 0.873. This implies that isolates predicted to be resistant were usually resistant, but many resistant isolates were missed. Gene-level and phylogeny-based predictions identified a somewhat larger proportion of resistant isolates, at the cost of lower precision (Fig. 4c–e). Performance also depended strongly on the background withheld from training (Extended Data Fig. 8b–e; Supplementary Table 16).

Together, these results show that genomic background associated with divergence among clades accounts for much of the predictive information available when related isolates occur in both training and test sets. Individual variant features supported the strongest overall prediction across held-out backgrounds, but incomplete detection of resistant isolates and substantial variation among backgrounds limited generalization. This pattern is consistent with evidence from bacterial resistance prediction that population structure and sampling composition can limit transfer across lineages^59^.

### Within-clade feature importance identifies established azole-response genes alongside broader antifungal-response candidates

Since fluconazole-resistance prediction varied across genetic backgrounds, we trained random forests separately within clades 1, 3 and 4 and PCA cluster 6, using individual variants or gene-based features recording whether an isolate carried a qualifying variant with predicted HIGH or MODERATE functional impact (Methods, Within-group random forest feature importance; Extended Data Fig. 9a–d; Supplementary Table 16). The leading gene-level features differed substantially among backgrounds: only *ERG11*, *EFG1* and *RCA1* appeared in more than one top-20 list, and PCA cluster 6 shared none with the named clades (Fig. 5a–e; Supplementary Tables 17–20). *ERG11* was the only gene shared by all three named clades. The other shared features also had published antifungal-response evidence: *EFG1* affects azole susceptibility and responses to caspofungin-induced cell-wall stress^60,61^, whereas *RCA1* deletion increases resistance to fluconazole and caspofungin^62^.

**Fig. 5.**
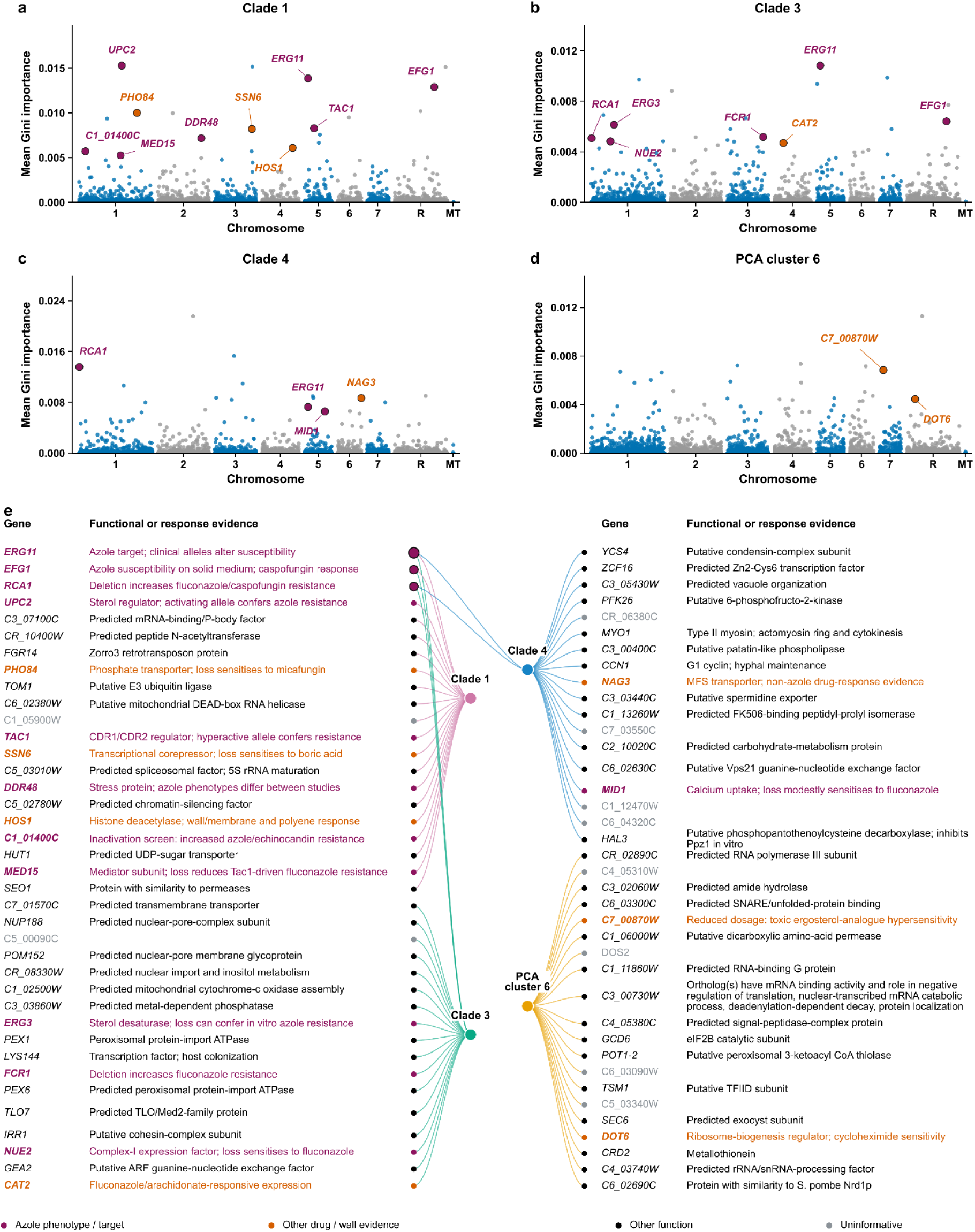
Gene-level predictors of fluconazole resistance within genetic backgrounds. **a–d**, Mean random forest Gini importance for clade 1 (**a**; 271 isolates, 167 resistant), clade 3 (**b**; 63 isolates, 21 resistant), clade 4 (**c**; 62 isolates, 10 resistant) and PCA cluster 6 (**d**; 63 isolates, 12 resistant). Each point represents a gene at its genomic midpoint; alternating blue and grey distinguish chromosomes. Inputs were binary gene-level carrier indicators derived from the shared variant catalogue, using nuclear and mitochondrial HIGH- or MODERATE-impact alleles carried by no more than 5% of the original 871 fluconazole-phenotyped isolates. Gini importance measures a feature’s contribution to reducing class impurity across forest splits. Importance was averaged across 25 outer-training fits for clade 1 (five repeats of five fold cross-validation) and 15 fits for each other group (five repeats of three fold cross-validation), assigning zero when a feature was not retained. Hyperparameters were selected by inner cross-validation. Labelled genes are among their group’s top 20 and have the direct or indirect response evidence indicated in **e**. **e**, Network connecting the four genetic backgrounds to their 20 highest-ranked genes, comprising 76 distinct genes and 80 group–gene memberships. Genes are arranged in two banks around the group nodes. Edge colours identify groups; gene-marker area scales with the number of linked groups, and dark outlines identify shared genes. Gene symbols, descriptions and markers share an evidence colour: magenta indicates published azole-response or target evidence; orange indicates other drug responses, cell-wall biology or explicitly qualified indirect responses; black indicates other annotated or predicted functions; grey indicates uninformative reference descriptions. Full reference descriptions, evidence terms and supporting sources are supplied with the source data.

In Clade 1, the established azole-resistance regulators *UPC2* and *TAC1*, together with genes linked to broader drug responses. These included *PHO84*, *SSN6* and *HOS1*, whose disruption increases susceptibility to micafungin, boric acid and polyenes, respectively^63–65^. *DDR48* has reported azole-response phenotypes that differ between studies^66,67^, whereas inactivation of C1_01400C (orf19.3329) increased resistance to azoles and echinocandins in a chemogenomic screen^41^. Clade 1 also prioritized *MED15*, a Mediator subunit whose deletion reduces fluconazole resistance in strains carrying gain-of-function *TAC1* alleles^68^. Thus, the leading features included both established resistance determinants and genes with broader or context-dependent effects on drug susceptibility (Fig. 5a,e; Supplementary Table 17).

In clade 3, additional features with direct azole-response evidence included *ERG3*, *FCR1* and *NUE2*: their disruption can confer azole resistance, increase fluconazole resistance and increase fluconazole susceptibility, respectively^40,69,70^. *CAT2* provided a more indirect connection through altered expression in biofilms exposed to fluconazole combined with arachidonic acid^71^ (Fig. 5b,e; Supplementary Table 18). Clade 4 prioritized *NAG3*, associated with cycloheximide resistance, and *MID1*, whose deletion modestly increases fluconazole susceptibility^72,73^ (Fig. 5c,e; Supplementary Table 19). In PCA cluster 6, C7_00870W (orf19.7029) and *DOT6* had evidence involving a toxic ergosterol analogue and cycloheximide, respectively^74,75^ (Fig. 5d,e; Supplementary Table 20). These connections support their consideration as candidates without establishing that the observed variants affect fluconazole resistance.

Variant-level importance showed two prominent concentrations of high-ranking variants in clade 1, on chromosomes 5 and R (Extended Data Fig. 9e; Supplementary Table 21). Within these regions, Erg11 D116E and Sec65 E213D were the highest-ranked nonsynonymous variants in their respective genes, although neither was the regional maximum. *SEC65* encodes a component of the protein-targeting machinery, and its disruption increases susceptibility to fluconazole and posaconazole in a chemogenomic screen^41^. The other clades showed more dispersed importance profiles, with isolated high-ranking variants rather than similarly pronounced clusters of elevated importance (Extended Data Fig. 9f–h; Supplementary Tables 22–24).

### Scores from variant effect predictors capture information associated with fluconazole resistance

Well curated genotypic resistance data helps explain the resistance phenotype of strains. However, many mutations present in our dataset have never been reported before, limiting the usefulness of previous mutations in interpreting these variants. Here, we explored whether scores generated by Variant Effect Predictors (VEPs) could be used to predict resistance. VEPs predict the impact of an amino acid substitution in a protein based on evolutionary protein history (GEMME)^76^, protein language models (ESM)^77^ or potential impact on the protein structure stability (FoldX)^78^. For GEMME and ESM, lower scores indicate a stronger predicted impact on the protein, with 0 representing no predicted effect. For FoldX, positive ΔΔG values predict that a substitution will be destabilizing, with 0 representing no predicted effect. We hypothesized that substitutions predicted by VEPs to have a greater impact on protein function or stability would be more likely to confer resistance. This signal could then be captured by a machine learning algorithm to predict resistance without prior knowledge of the resistance phenotype associated with substitutions.

To evaluate how well the models generalize beyond closely related strains, we performed clade-grouped CV, in which isolates from the same clade were kept within the same fold (Fig. 6 a). As a reference, we trained a random forest using only FungAMR confidence scores (median balanced accuracy = 76.5% (IQR 72.7%-85.7%), median precision = 85.7% (IQR 83.3%-94.7%), median recall = 55.6% (IQR 50.0%-75.0%)). We then compared this reference with models using VEP scores for 23 proteins listed in FungAMR.

**Fig. 6.**
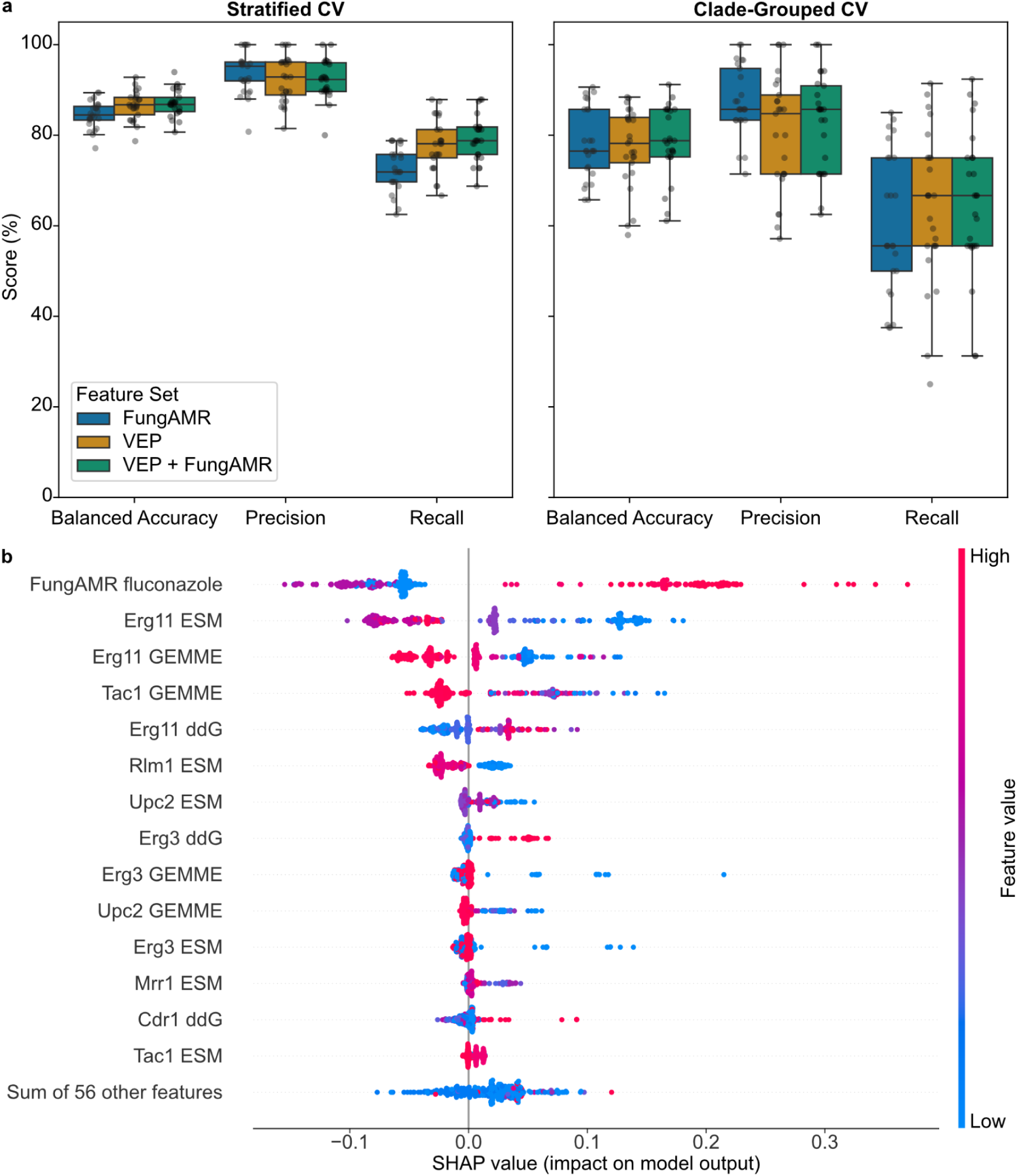
Random-forest prediction of binary fluconazole resistance from FungAMR and variant effect predictors. **a,** Balanced accuracy, precision and recall under stratified CV and clade-grouped CV using three different sets of features: FungAMR fluconazole scores (FungAMR), Variant effect predictor scores (VEP) and the combination of both (VEP + FungAMR). Individual data points represent the performance metrics over 25 total folds (five folds repeated five times) and boxes show the IQR, with the median as the centre line, quartiles as box limits and whiskers extending to 1.5 times the IQR. n=382 strains. **b,** SHAP values computed for the resistant class (1) of the VEP + FungAMR clade-grouped stratified CV. A positive SHAP value indicates a contribution toward a resistance prediction and negative values indicate a contribution towards a susceptibility prediction.

Surprisingly, models with only VEPs as features perform similarly as FungAMR scores with a lower precision, but a higher median recall (median balanced accuracy = 78.2% (IQR 73.9%-83.9%), median precision = 84.8% (IQR 71.4%-88.9%), median recall = 66.7% (IQR 55.6%-75.0%)). These results suggest that VEP scores contain meaningful information associated with resistance that is not fully captured by our current knowledge of resistance mutations, but yields a larger uncertainty than using curated data. Combining FungAMR and VEP scores did not substantially improve performance compared with either feature set alone (median balanced accuracy = 78.8% (IQR 75.2%-85.7%), median precision = 85.7% (IQR 71.4%-90.9%), median recall = 66.7% (IQR 55.6%-75.0%)), indicating that the information provided by the two feature sets is partially redundant. Overall, the high precision indicates that resistance predictions are generally correct, whereas the lower recall indicates that many resistant isolates are missed. This is most likely because some resistance mutations are not represented by any of the features used in the models, for example in genes not included, or because some resistance mutation patterns are too rare in our dataset for the model to learn them.

To identify which features contributed to resistance predictions, we used SHapley Additive exPlanations (SHAP) values. Here, a positive SHAP value indicates that the feature impacted the model towards resistance prediction. As expected, the most important feature is the FungAMR score, with a high positive confidence score contributing to a resistance prediction (Fig. 6b, red datapoint towards the right). Then, VEP scores obtained for the drug target Erg11 are among the most important. If a strain harbors an amino acid change predicted to impact the protein or destabilize it, the model will most likely predict resistance. After, there are the Zn_2_-Cys_6_ transcription factors Tac1 and Upc2 that both have been identified as drivers of azole resistance before^79^. They are followed by Erg3, for which loss of function mutations have been associated with resistance^79^. An interesting feature contributing to resistance prediction is Rlm1, but for ESM scores only. This transcription factor is part of the PKC signaling pathway, which has been identified as a crucial part of the cell response to antifungals^80^.

Together, these results show that predicted protein effects capture resistance-associated information without requiring a curated resistance annotation for each substitution. These findings suggest that restricting our focus to well characterized and established targets can be as relevant for resistance prediction as how much of the genome is covered. Focusing on established resistance proteins and summarising substitutions by predicted functional impact may retain relevant signals while reducing the overall volume of data required. Ultimately, the value of VEPs lies in providing a biological connection between sequence changes and resistance, potentially extending the usefulness of known mechanisms to predict the effect of uncharacterized substitutions.

### Convergent genetic evidence prioritizes resistance gene candidates

Comparison of top genes associated with the R phenotype from the GWAS and random forest classifier showed the clearest agreement for the genes *ERG11*, *TAC1*, *UPC2* and *ERG3* for fluconazole (Fig. 3b; Fig. 5e; Supplementary Tables 10,17–20). Most other high importance genes from RF features did not meet the GWAS thresholds and vice versa. The differences may partly reflect the distinct questions addressed: GWAS tested associations with MIC across the population, whereas RFs predicted binary phenotype within individual backgrounds. Together, the analyses converged on established resistance genes, while identifying additional candidates whose predictive value depended on genetic background.

To narrow the network-derived candidates to genes with stronger evidence for involvement in resistance, we examined enrichment among R phenotype isolates lacking high confidence FungAMR resistance substitutions (Methods, Prioritizing candidate genes). This identified 36 enriched gene–antifungal associations across 30 genes after accounting for genetic background: 21 for fluconazole, four each for voriconazole and posaconazole, two for caspofungin and five for micafungin (Extended Data Fig. 10a). The fluconazole candidates included *RCA1*, *PHO84*, *MRR1*, *FCR1* and *ACU1*, alongside genes involved in sterol metabolism. The echinocandin candidates comprised *FKS1* and *C2_01510C* for caspofungin, and *INO1*, *C2_07760W*, *HBR3*, *C3_03410C* and *CDC68* for micafungin. No gene met the enrichment criteria for anidulafungin, although only one R phenotype isolate remained available for this comparison.

Many of these candidates occurred together in the same isolates, making their individual contributions difficult to distinguish. Among fluconazole R phenotype isolates, 61 of 62 carried variants in at least two qualifying genes, and one isolate carried variants in 13. Multiple qualifying genes were also carried by 17 of 45 voriconazole and 34 of 63 posaconazole R phenotype isolates (Extended Data Fig. 10b). After accounting for relevant co-carried genes, eight associations remained significant: *ERG3*, *GEA2* and *C5_03010W* for fluconazole; *ERG3*, *ERG251*, *GLO1* and *C5_00090C* for voriconazole; and *TAC1* for posaconazole (Extended Data Fig. 10c). The *TAC1*–posaconazole association was retained across all alternative co-carriage selection thresholds examined. Seventeen other associations, including *RCA1* and *PHO84* for fluconazole, no longer met the significance criterion. Six associations had no selected co-carried partners, while the five micafungin associations could not be evaluated after adjustment because informative comparisons were unavailable. These outcomes distinguish the retained candidates but do not exclude biological contributions from the other genes.

Within the retained candidates, nine individual variants were significantly enriched (Extended Data Fig. 10d). For fluconazole, these were E242K and E241K in *C5_03010W*, and E94K, G93fs, G90fs, E89K and D88N in *GEA2*. For voriconazole, A211T in *GLO1* and F6fs in *C5_00090C* were enriched. These findings nominate specific alleles for functional investigation within the prioritized genes. No individual variant met the significance criterion for the retained *ERG3*, *ERG251* or *TAC1* associations. The contrast between gene-level and individual-variant evidence was particularly informative for *ERG3* and *ERG251*. Sixteen distinct *ERG3* variants occurred among R phenotype isolates in each of the fluconazole and voriconazole cohorts, with each variant carried by only one or two such isolates. For voriconazole, eight distinct *ERG251* variants were each carried by a single R phenotype isolate. Thus, the significant pooled associations were distributed across numerous individually rare alleles. A plausible biological explanation is that several different variants affect the same gene and contribute to the R phenotype, while each remains too uncommon to produce a detectable association alone. Pooling their contributions therefore revealed a shared gene-level signal.

## Discussion

This study asked how well whole-genome sequencing can explain and predict antifungal resistance in *C. albicans* across different genetic backgrounds. The collection of more than 1000 whole-genome sequenced isolates helped resolve major lineages and uncovered extensive loss of heterozygosity across them, a mechanism that can influence the effects of adaptive mutations^81^. Because of the structured populations within this species, shared ancestry needed to be considered alongside individual mutations for the association with and the prediction of resistance mutations.

Our work shows the importance of having high-quality resistance data to make predictions, although the currently available data is incomplete. Indeed, curated mutations from FungAMR were strong indicators of resistance, but their absence was less informative, indicating that many more mutations or genetic mechanisms can cause resistance than what is currently reported. Catalogue-based approaches therefore appear better suited for recognition of established mechanisms than for excluding resistance phenotypes. Gene-level GWAS associations and within-background random forest features converged on *ERG11*, *TAC1*, *UPC2* and *ERG3* that are well-established determinants of resistance to azoles. However, the additional candidate genes prioritized by the two approaches overlapped less. This could reflect their different analytical aims—detecting population-wide associations with MIC versus predicting the resistance within genetic backgrounds—as well as differences in sample size and the genetic variation represented in each analysis. On the other hand, mutations in *FKS1* appear to be largely sufficient to explain resistance to echinocandins.

A particularly informative result was that pooling rare protein-altering variants uncovered associations beyond those detected by testing variants individually. Gene-level analysis strengthened the association of the transcriptional regulator *TAC1* with fluconazole resistance and recovered established determinants such as *UPC2,* another transcription factor, and *ERG3,* an enzyme of the ergosterol pathway, alongside additional candidates. This is consistent with different rare changes in the same gene contributing to a shared susceptibility phenotype even when individual alleles provide insufficient evidence on their own. This suggests that while the mutational catalogue of resistance is incomplete, the list of the genes involved may be closer to completion. A better understanding of which mutations within each of these known loci cause resistance would therefore highly improve the interpretation of genomic variation. In the meantime, aggregation therefore offers a practical way to investigate resistance mechanisms distributed across multiple mutations. Candidate associations extending beyond drug targets also suggest contributions from broader cellular processes that merit experimental investigation. Of particular interest for future investigation are *ERG3,* also uncovered with other methods and *GEA2* and *C5_03010W* for fluconazole; *ERG3*, *ERG251*, *GLO1* and *C5_00090C* for voriconazole; and *TAC1* for posaconazole. These gene-level associations remained significant after accounting for genetic background and selected co-carried genes, with the *TAC1*–posaconazole association retained across all alternative co-carriage selection thresholds examined. Separate individual-variant analyses identified enriched alleles in *GEA2*, *C5_03010W*, *GLO1* and *C5_00090C*, providing specific candidates for functional testing. The retained *ERG3* and *ERG251* associations instead involved numerous individually rare alleles.

The overlap between GWAS and within-clade random forest features further connected discovery with prediction. Both approaches prioritized *ERG11*, *TAC1*, *UPC2* and *ERG3*, while random forest classifiers identified additional features that differed among genetic backgrounds. These patterns suggest a shared set of established mechanisms alongside potentially context-dependent contributions but their individual contributions cannot be quantified with the current dataset. Interestingly, two of these are enzymes of the pathway targeted by azoles and the other two are transcription factors, suggesting two frequent alternative genetic paths towards resistance. Phylogeny-based features predicted resistance almost as well as individual variants under stratified validation, showing how much predictive information resides in population structure. Unfortunately, such association is likely an artefact of the sampling scheme as resistance evolves *de novo* within each lineage such that clades themselves cannot be causally linked to resistance. Individual variants nevertheless provided the strongest average transfer to backgrounds absent from training. Their high precision means that resistance predictions were usually correct, supporting their usefulness for flagging likely resistant isolates. However, limited recall means that many resistant isolates were wrongly classified as susceptible. Improving the detection of these missed isolates while maintaining the reliability of resistance predictions remains an important objective.

The fact that pooling the rare variants within genes improved the association for some genes suggests that there are more variants contained in known resistance genes that could contribute to resistance. Machine learning across Saccharomycotina yeasts has identified Erg11 sites predictive of fluconazole resistance that differed from those previously implicated in clinical *C. albicans* isolates^82^. The use of Variant Effect Predictors (VEPs) offers a powerful approach in this context in the absence of experimental data as they allow extending predictions beyond individually documented substitutions. Under grouped validation, predicted protein effects supported performance similar to curated scores while increasing recall. Functional summaries like variant effect predictors may therefore help connect sequence changes with resistance even when their individual effects have not been catalogued. Limited improvement from combining VEP and FungAMR scores suggests that they capture partly shared information. Their use of Erg11, Tac1, Upc2 and Erg3 as features reinforces this interpretation, while the Rlm1 signal raises a further connection to antifungal stress responses^80^. Further improvements may come from refining this functional interpretation and incorporating additional resistance mechanisms, rather than simply adding more genomic features.

Our findings motivate broader functional testing and independent validation. Uneven sampling, heterogeneous susceptibility measurements and limited sampling of echinocandin resistant isolates constrain discovery, while the VEP analysis covers a selected cohort and known resistance-associated proteins. Direct comparisons between prediction strategies will require matched cohorts, and previously unseen substitutions provide an important next test. Experimental assessment of candidate variants, variant combinations and genetic backgrounds could turn these associations into mechanisms and expand resistance catalogues. Together, these approaches provide a foundation for moving genomic resistance prediction beyond recognition of familiar mutations towards interpretation of a wider range of genetic variation and use for clinical treatment decisions.

## Extended Data Figures

**Extended Data Figure 1.**
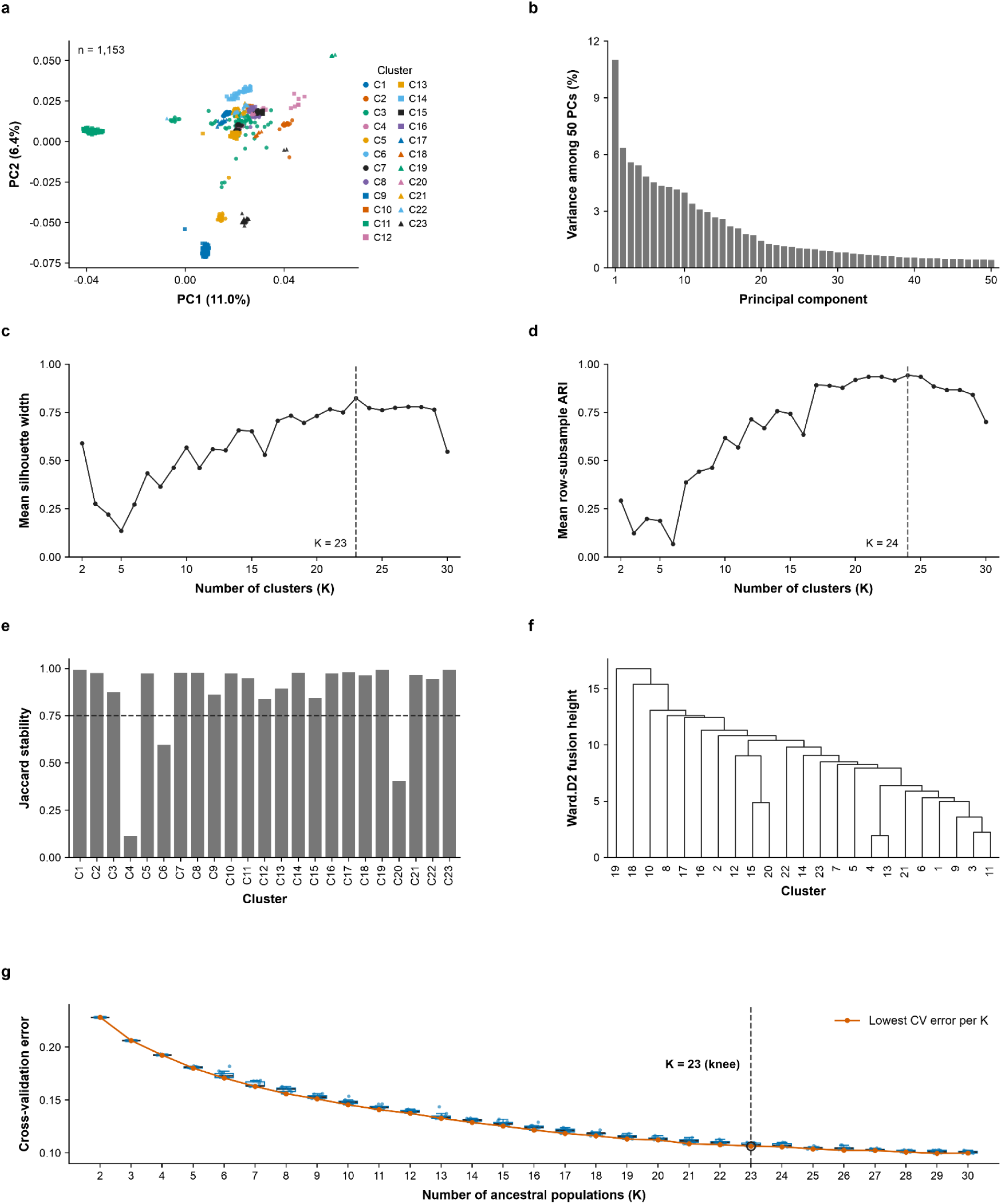
Population structure assessed using PCA, clustering and ADMIXTURE. **a,** Principal-component analysis (PCA) of 1,153 isolates using 23,356 linkage disequilibrium-pruned SNPs. Colours and marker shapes identify the 23 k-means clusters. **b,** Variance represented by the first 50 principal components. Percentages in **a** and **b** are relative to the summed eigenvalues of these 50 components. **c,** Mean silhouette width across K = 2–30; the dashed line marks its maximum at K = 23. **d,** Mean adjusted Rand index (ARI) across 20 independent subsamples containing 80% of isolates, drawn without replacement. ARI measures agreement with the full-data clustering after accounting for chance; the dashed line marks its maximum at K = 24. **e,** Clusterwise Jaccard stability at K = 23 across 200 bootstrap replicates. The dashed line marks the 0.75 stability threshold. **f,** Hierarchical clustering of the 23 centroids in centered and scaled PC1–PC20 space, using Euclidean distances and Ward.D2 linkage. **g,** ADMIXTURE tenfold cross-validation (CV) error for K = 2–30, with 10 independent runs per K. Blue points show individual runs; boxes show the interquartile range, center lines the median, and whiskers the most extreme values within 1.5 times the interquartile range. The orange line connects the lowest error at each K. The dashed line and open circle highlight K = 23 as the knee of the curve; the overall minimum occurs at K = 29.

**Extended Data Figure 2.**
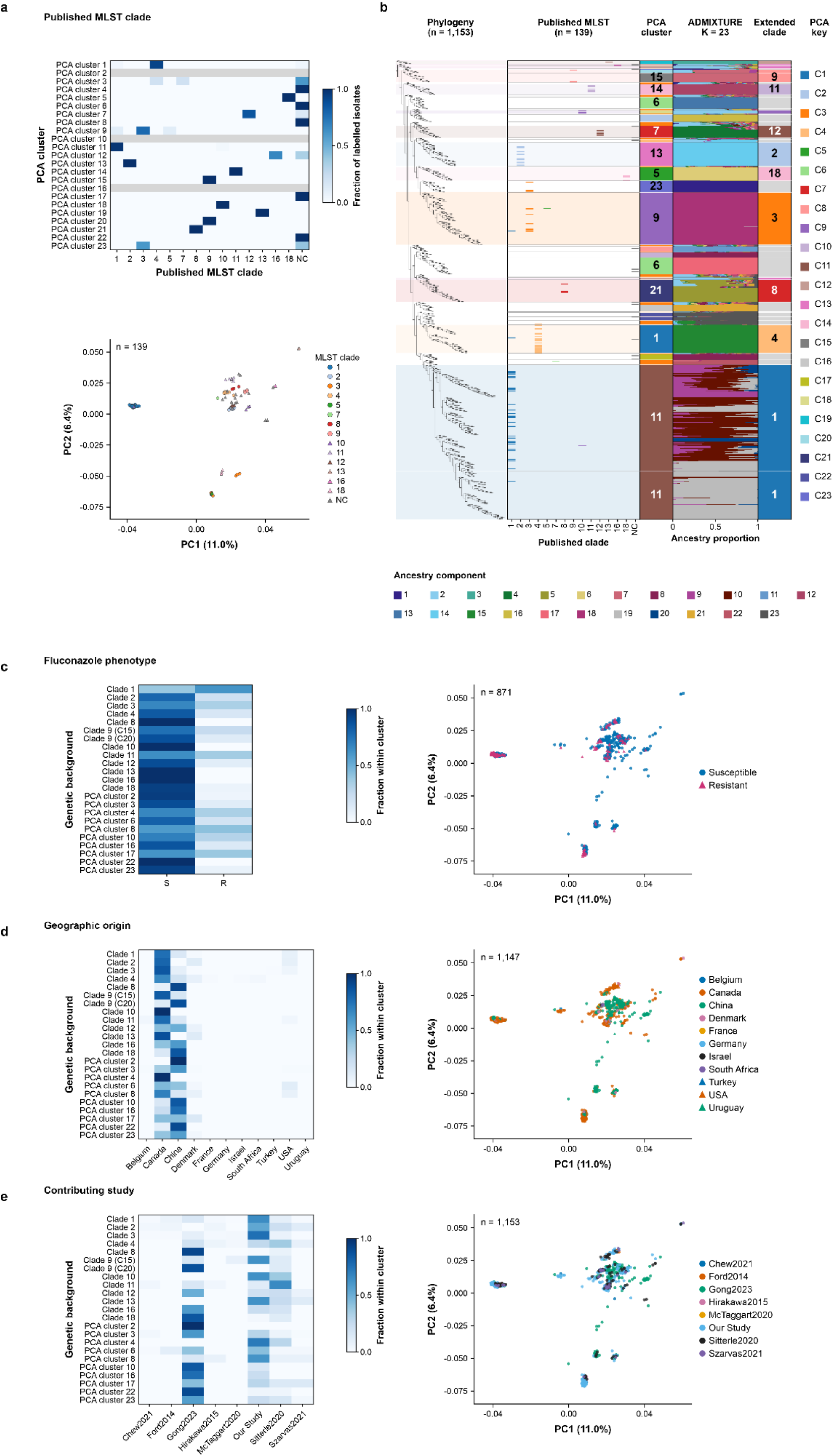
Concordance of published clade labels with population structure and extension of clade assignments. **a,** Overlap between published multilocus sequence typing (MLST) clades and PCA clusters (top), and PCA coordinates coloured by published MLST clade (bottom), for 139 labeled isolates. Heatmap rows show all 23 PCA clusters in numerical order. Values are fractions of MLST-labelled isolates within each cluster; grey rows indicate clusters without published MLST labels. **b,** Maximum-likelihood tree of 1,153 isolates, displayed without branch-length scaling and in the same tip order as main Fig. 1. Aligned tracks show published MLST labels, PCA clusters, individual ADMIXTURE ancestry proportions at K = 23 and extended clade assignments. Each horizontal row represents the same isolate across tracks. Published MLST labels occupy separate clade columns, with colours matching the extended-clade track. Pale bands show the extent of named-clade assignments. Horizontal separators mark PCA cluster boundaries; bold strip numbers identify PCA clusters and extended clades. Published labels provided anchors for extending clade assignments, supported by phylogenetic placement and ancestry profiles, with PCA cluster boundaries defining each extension. Grey denotes clusters without a supported named-clade assignment. Disjoint occurrences of a cluster retain their original tree positions. ADMIXTURE colours identify ancestry components, and each isolate’s bar sums to one. Values derive from the lowest-cross-validation-error K = 23 run, fitted using 1,201 input records and 82,364 SNPs and matched to all displayed isolates. **c–e,** Cluster composition (left) and PCA coordinates (right), coloured by fluconazole susceptibility (c; n = 871, comprising 607 susceptible and 264 resistant isolates), geographic origin (d; n = 1,147) and contributing study (e; n = 1,153). Heatmap rows retain the original PCA clusters, labeled by assigned clade where available. Values represent fractions among isolates with the corresponding metadata within each cluster. Scatter-plot colours and marker shapes identify categories in the keys. NC, previously reported new clade; S, susceptible; R, resistant. PCA percentages are relative to the summed eigenvalues of the first 50 reported components.

**Extended Data Figure 3.**
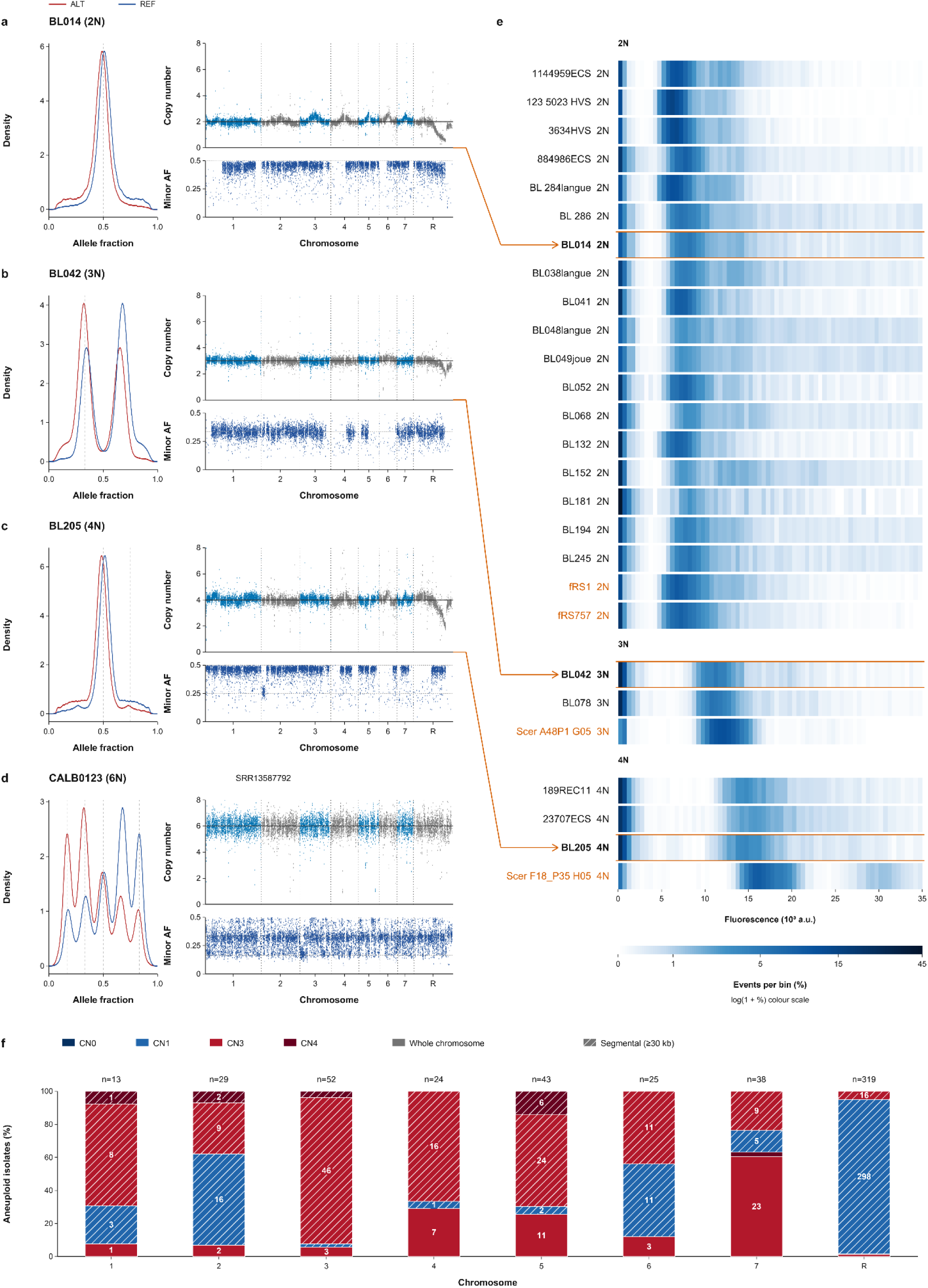
Copy-number variation across the isolate collection. **a–d,** Genome-wide profiles for the isolates BL014 (a, 2N), BL042 (b, 3N), BL205 (c, 4N) and CALB0123_SRR13587792 (d, 6N). The left column shows the density of alternate-allele (ALT; red) and reference-allele (REF; blue) fractions; vertical dashed lines mark expected allele fractions for the assigned ploidy. The middle column shows depth-derived absolute copy-number estimates across chromosomes 1–7 and R (top), with alternating blue and grey distinguishing adjacent chromosomes and a horizontal reference line marking baseline ploidy, and median minor-allele fraction (AF) in 1-kb bins (bottom). In BL205, a short chromosome 2 interval has a minor-allele fraction near 0.25 without a corresponding local copy-number increase, despite predominantly 0.5 allele-fraction peaks elsewhere. The 6N sample was not available for flow-cytometry testing. The collection comprised 1,127 2N, seven 3N, 12 4N and seven 6N isolates; no 5N isolate was detected. **e,** Flow-cytometry fluorescence distributions for all 23 tested isolates belonging to the 1,153-isolate cohort and four reference-control wells: *C. albicans* fRS1 and fRS757 (2N) and *S. cerevisiae* A48P1 (3N) and F18_P35 (4N), acquired in the same run. Rows are grouped by increasing recorded ploidy, with reference controls immediately after cohort isolates of the same ploidy. The fRS1 and fRS757 recordings contain 5,000 and 4,786 events, respectively. Scer denotes *S. cerevisiae*; control isolate names and their ploidy labels are orange. Row labels report ploidy recorded in the experimental plate map, rather than new assignments from heatmap intensity. Each cell represents the percentage of all recorded events in a 500-unit bin of the linear green-fluorescence height channel (GRN-B-HLin). The fluorescence axis displays 0–35,000 arbitrary units; events outside this range remain in the normalization, and no additional gating is applied. A common log(1 + percentage) colour scale spans 0–45% of events per bin; no observed bin is clipped by the colour scale. Orange arrows connect the copy-number profiles of BL014 (2N), BL042 (3N) and BL205 (4N) to their outlined heatmap rows; these wells contain 5,000, 2,867 and 2,720 events, respectively. *S. cerevisiae* controls are identified separately because fluorescence is not directly interchangeable between species. Later acquisitions at altered concentrations are not pooled with the initial plate. **f,** Chromosome-specific composition of aneuploidy among the 1,127 diploid isolates. Solid segments denote whole-chromosome events and hatched segments denote segmental events spanning at least 30 kb. Colours indicate copy-number state (CN0, CN1, CN3 or CN4); for segmental events, the dominant state is the modal copy number among altered loci in qualifying segments within an isolate and chromosome. Bars sum to 100% of isolates with an event on the corresponding chromosome. Numbers within segments are isolate counts, shown where space permits; ‘n’ above each bar is the total number of isolates with an event on that chromosome. Isolates with events on multiple chromosomes contribute to each relevant chromosome.

**Extended Data Figure 4.**
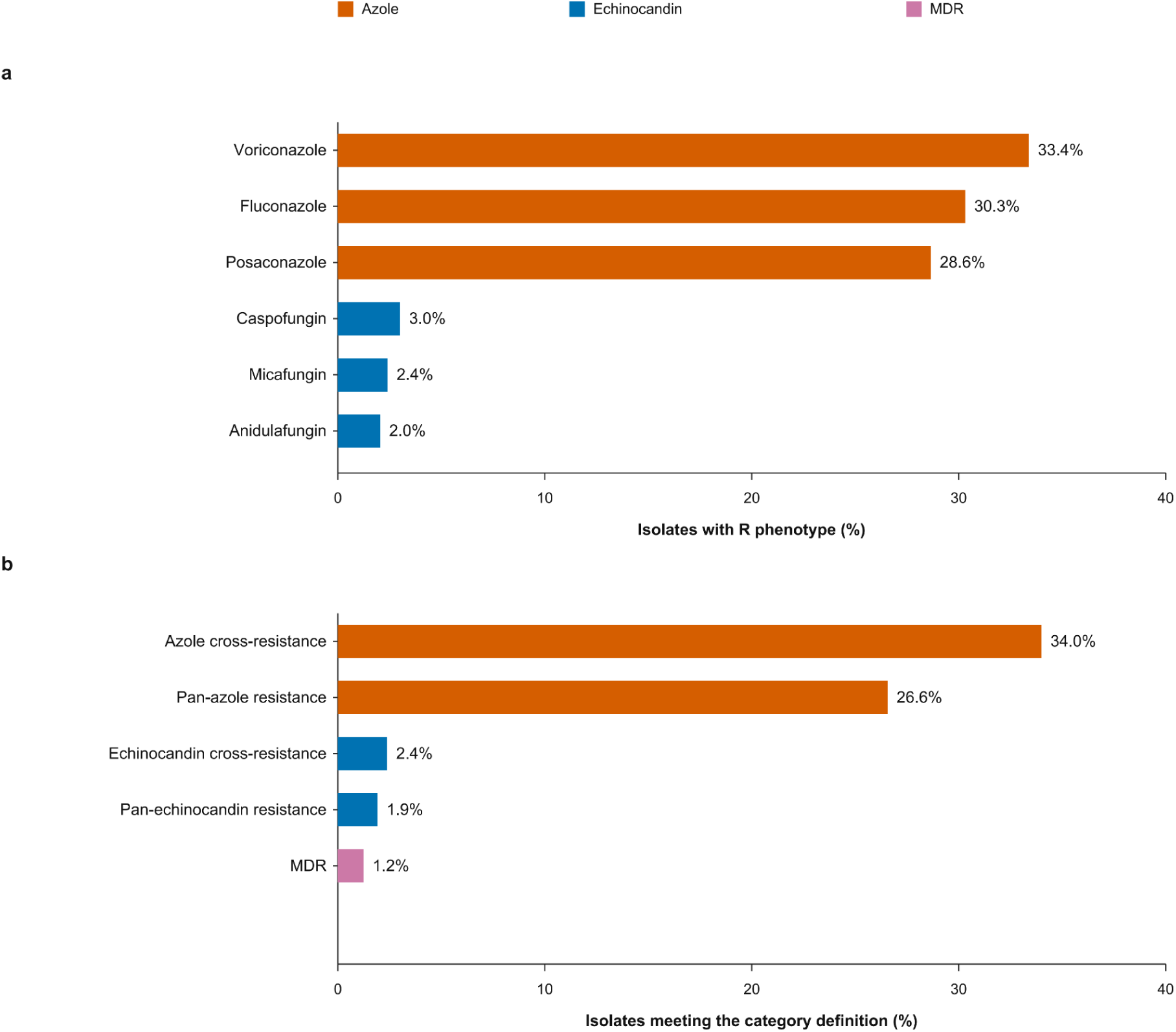
Distribution of antifungal resistance, within-class cross-resistance and multidrug resistance across the cohort. **a,** Frequencies of the R phenotype among isolates tested against each of six antifungals in the 1,153-isolate collection. Bars show the percentage with the R phenotype for voriconazole (*n* = 659 tested), fluconazole (*n* = 871), posaconazole (*n* = 611), caspofungin (*n* = 766), micafungin (*n* = 667) and anidulafungin (*n* = 589). **b,** Frequencies of five overlapping phenotype categories among isolates with the relevant testing data. Azole and echinocandin cross-resistance denote R phenotypes for at least two drugs within the respective class, among isolates tested against at least two drugs in that class (*n* = 659 and *n* = 632, respectively). Pan-azole and pan-echinocandin resistance denote R phenotypes for all three drugs in the respective class, among isolates tested against all three (*n* = 610 and *n* = 575, respectively). MDR denotes an R phenotype for at least one azole and at least one echinocandin, among isolates tested against at least one drug from each class (*n* = 809). These categories describe phenotype patterns without implying a shared resistance mechanism. Colours distinguish azole (orange), echinocandin (blue) and MDR (pink) categories. Labels beside bars give percentages; both panels use the same percentage scale. Isolates may contribute to multiple drugs and categories; untested drug–isolate combinations are excluded from the relevant denominators. Counts are provided in the source data.

**Extended Data Figure 5.**
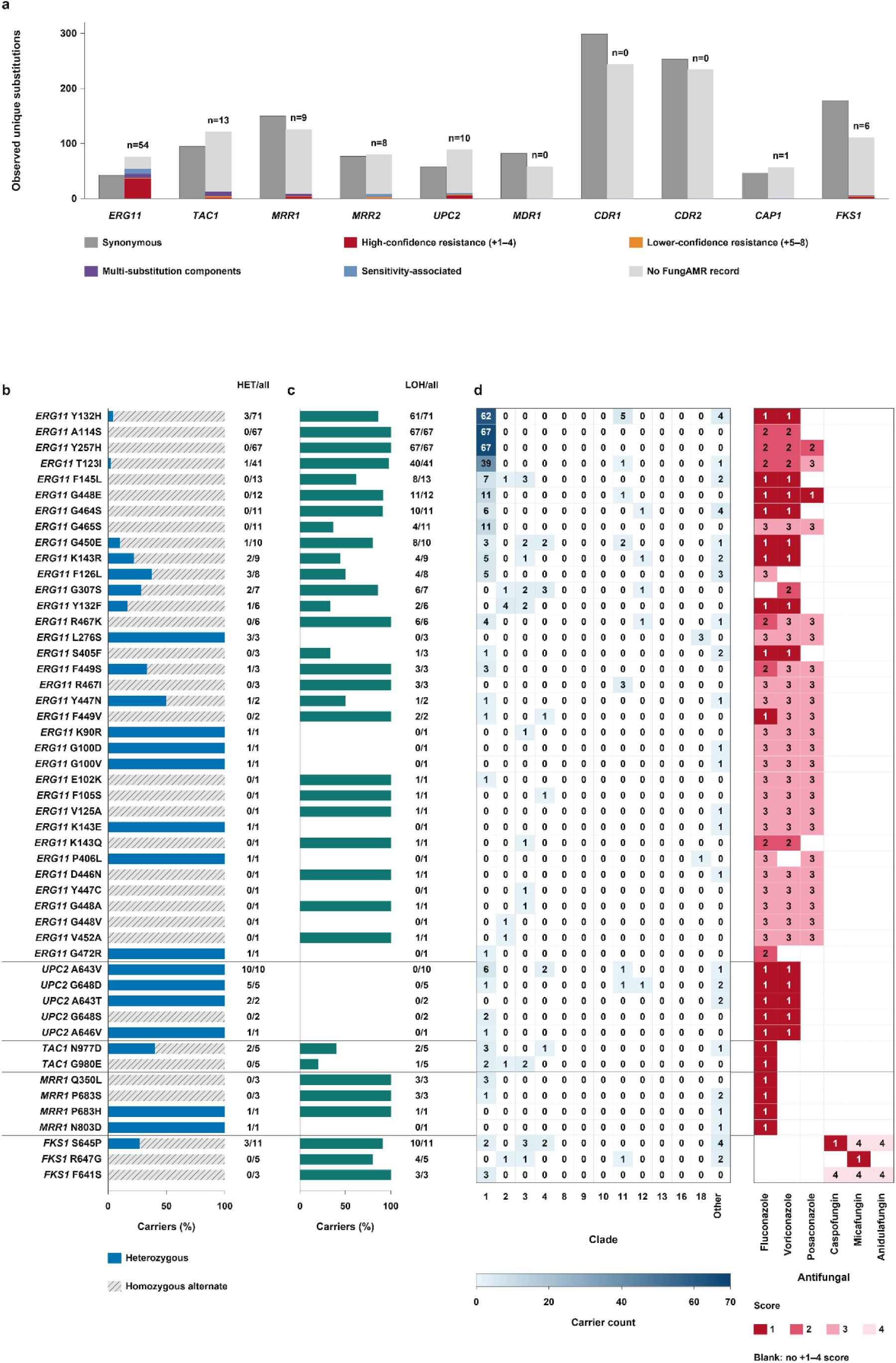
FungAMR evidence, zygosity, LOH and clade distribution of substitutions in antifungal-resistance-associated genes. **a**, Numbers of observed synonymous (dark grey) and unique nonsynonymous amino-acid substitutions in ten selected genes. Nonsynonymous bars are stacked by FungAMR confidence scores: high-confidence resistance (scores +1–4; red), lower-confidence resistance (scores +5–8; orange), resistance-associated components of multiple-substitution records (purple), sensitivity-associated substitutions supported by negative scores (blue), and substitutions without matching FungAMR records (light grey). Numbers above the nonsynonymous bars indicate substitutions with at least one matching record. Each nonsynonymous substitution contributes to one category, prioritising exact high-confidence resistance, exact lower-confidence resistance, components of resistance-associated combinations, sensitivity-associated evidence and unmatched substitutions, in that order. **b**, Proportions of heterozygous (blue) and homozygous-alternate (hatched grey) carriers of 49 high-confidence substitutions. HET/all labels indicate heterozygous carriers divided by all carriers. One and two copies of the specified alternate allele in diploid genotype calls define these states, respectively. **c**, Percentage of carriers whose corresponding gene overlaps a called LOH tract. Labels give LOH-positive carriers divided by all carriers; any overlap with the gene interval qualifies. Panels b and c use the same row order and denominators. **d**, Clade distribution of the same 49 substitutions (left) and their drug-specific high-confidence FungAMR annotations (right). Rows are grouped by gene, with ERG11 first, followed by other azole-associated genes and FKS1. Left-hand cell values and colour intensity indicate carrier counts; white indicates zero. Other denotes isolates without an assigned clade. Right-hand cells show resolved confidence scores +1 to +4, from dark red to pale pink; white denotes no high-confidence resistance record for that substitution–drug pair. Panel a considers FungAMR evidence across antifungals; panels b–d show high-confidence substitutions annotated for at least one of the six displayed drugs. Carrier counts are independent of resistance phenotype and co-carriage; an isolate with multiple substitutions contributes to each relevant row. LOH, loss of heterozygosity.

**Extended Data Figure 6.**
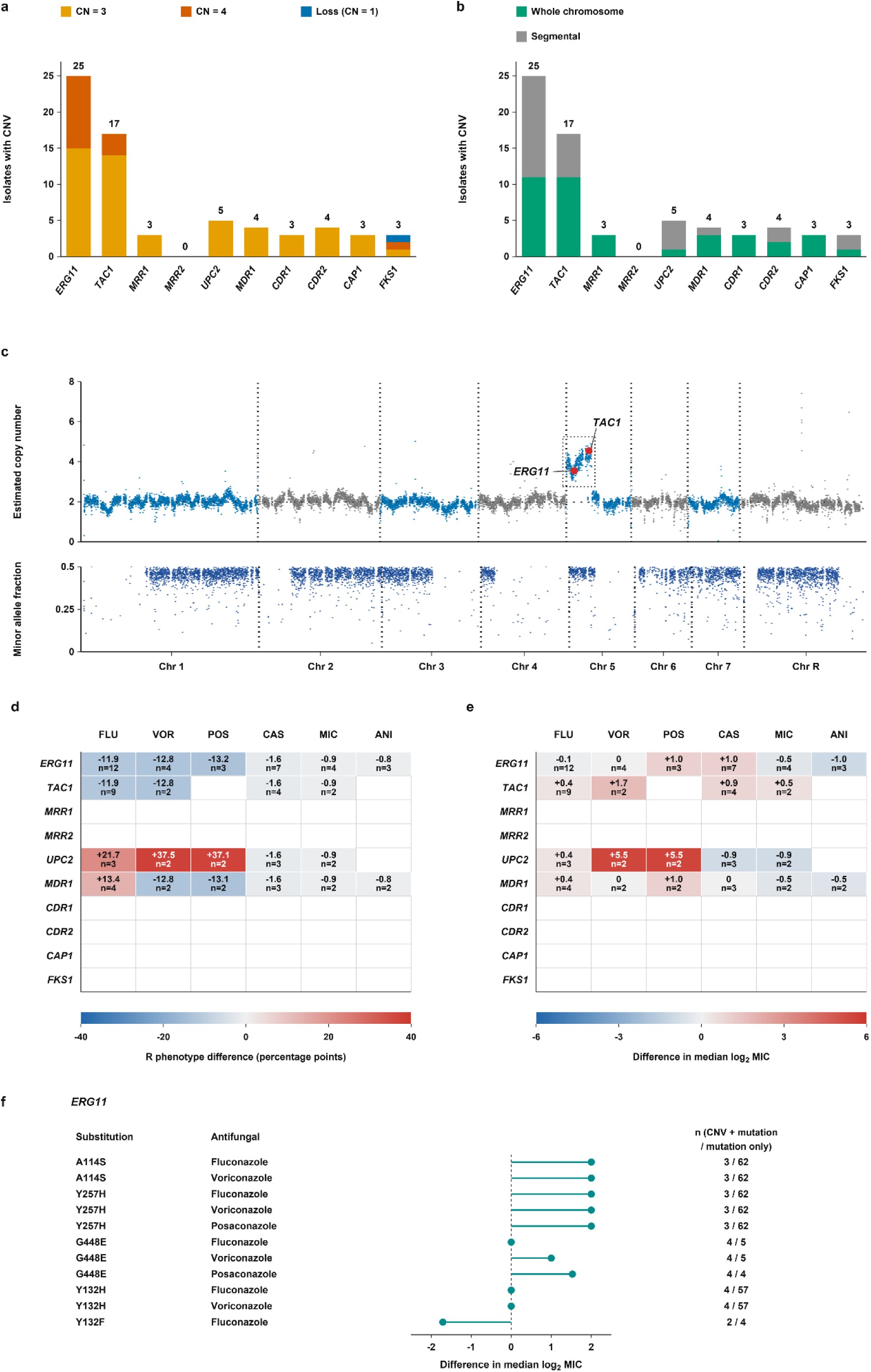
Copy-number variation at antifungal-resistance loci and associations with susceptibility. **a**, Numbers of isolates carrying a copy-number variant (CNV) at each resistance-associated gene among 1,127 diploid genomes. Orange, vermilion and blue indicate gains to copy number (CN) 3, gains to CN4 and losses to CN1, respectively. Numbers above bars indicate CNV-positive isolates. **b**, Genomic context of the same CNVs. Green indicates events in which the gene copy number matched a whole-chromosome copy-number change; grey indicates segmental events. Isolates with CNVs at multiple genes contribute to each relevant bar. **c**, Representative chromosome 5L amplification in isolate CALB0442_SRR1554302. Estimated copy number (upper plot) and minor-allele fraction (lower plot) are shown across chromosomes 1–7 and R. Alternating blue and grey distinguish adjacent chromosomes in the upper plot. Vertical dotted lines delimit chromosomes, and the horizontal dotted line marks CN2. The dotted rectangle highlights chromosome 5L amplification to approximately CN4, consistent with an i(5L)-like event encompassing ERG11 and TAC1. Larger red dots mark the measured copy numbers of the 1-kb bins containing each gene’s midpoint. **d**,**e**, Differences between gene-copy-number gain carriers and non-carriers in resistant fraction calculated from binary resistance phenotype, expressed in percentage points (d), and median log₂ MIC; e). All 219 diploid isolates carrying any high-confidence FungAMR substitution (signed-resolved scores +1 to +4) were excluded. Cells require at least two eligible gain carriers with the relevant phenotype. Annotations show the carrier-minus-non-carrier difference and carrier count (n). Red indicates higher values among carriers, blue indicates lower values and grey indicates values near zero; blank cells indicate insufficient carriers. **f**, Differences in median log₂ MIC between carriers of an ERG11 substitution with and without an ERG11 CNV. Isolates carrying high-confidence substitutions in other resistance genes were excluded; additional ERG11 substitutions were permitted. Lollipop points show median differences, with stems extending from zero. Counts give CNV-plus-mutation and mutation-only group sizes. Comparisons may share isolates; small groups limit inference. FLU, fluconazole; VOR, voriconazole; POS, posaconazole; CAS, caspofungin; MIC, micafungin; ANI, anidulafungin.

**Extended Data Figure 7.**
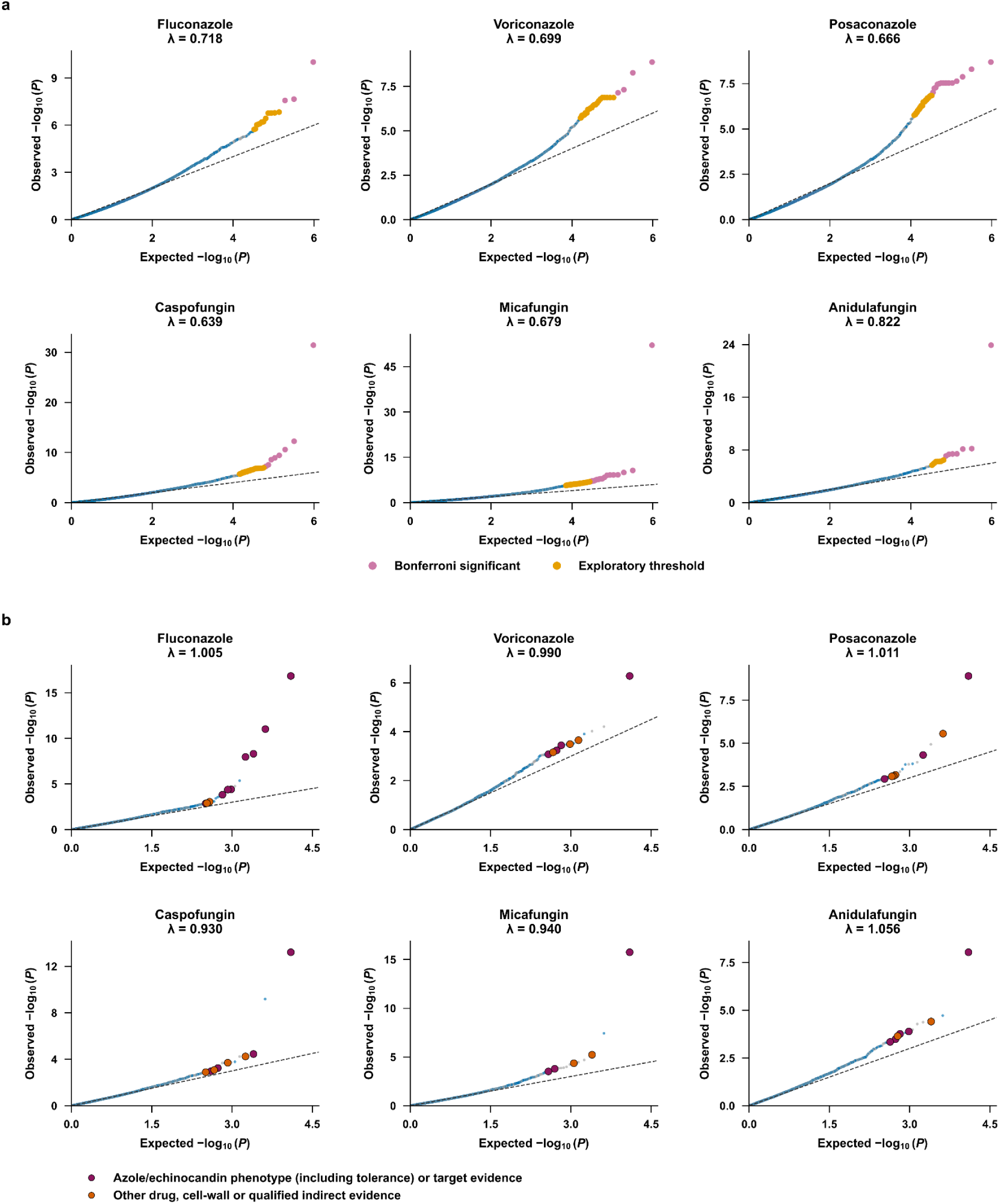
Quantile–quantile plots for variant and gene-level variant associations with antifungal susceptibility. **a,** Variant GWAS of individual SNPs and indels. **b,** Gene-level variant GWAS, aggregating predicted protein-altering variants carried by ≤5% of each drug-specific cohort into gene-level carrier indicators. Both analyses tested associations with log₂-transformed MIC. Within each panel, the upper row shows fluconazole, voriconazole and posaconazole, and the lower row shows caspofungin, micafungin and anidulafungin. The corresponding cohorts contained 871, 659, 611, 766, 667 and 589 isolates, respectively. Each point represents one tested variant (a) or gene (b), including nuclear and mitochondrial tests. Observed −log₁₀(P) values are plotted against those expected under a uniform null distribution, using expected probabilities (r − 0.5)/M, where r is the ascending P-value rank and M is the number of tests for that drug. Dashed diagonals indicate agreement with the null expectation. In a, pink and orange mark Bonferroni-significant (P < 0.05/M) and suggestive-only (0.05/M ≤ P < 1/M) associations, respectively. In b, enlarged outlined markers identify the same literature-supported top-20 genes highlighted in Fig. 3b: magenta denotes published azole/echinocandin phenotype or target evidence, and orange denotes evidence involving other antifungals, cell-wall phenotypes or indirect responses. Remaining points use the alternating blue and grey chromosome colours from Fig. 3. The genomic inflation factor, λ, equals the median association χ² statistic divided by the expected median of a χ² distribution with one degree of freedom (approximately 0.455). Variant statistics were obtained from P values; gene-level statistics were calculated as squared effect-to-standard-error ratios. Values below and above 1 indicate deflation and inflation of the median statistic, respectively. All tests are displayed without rescaling P values by λ.

**Extended Data Figure 8.**
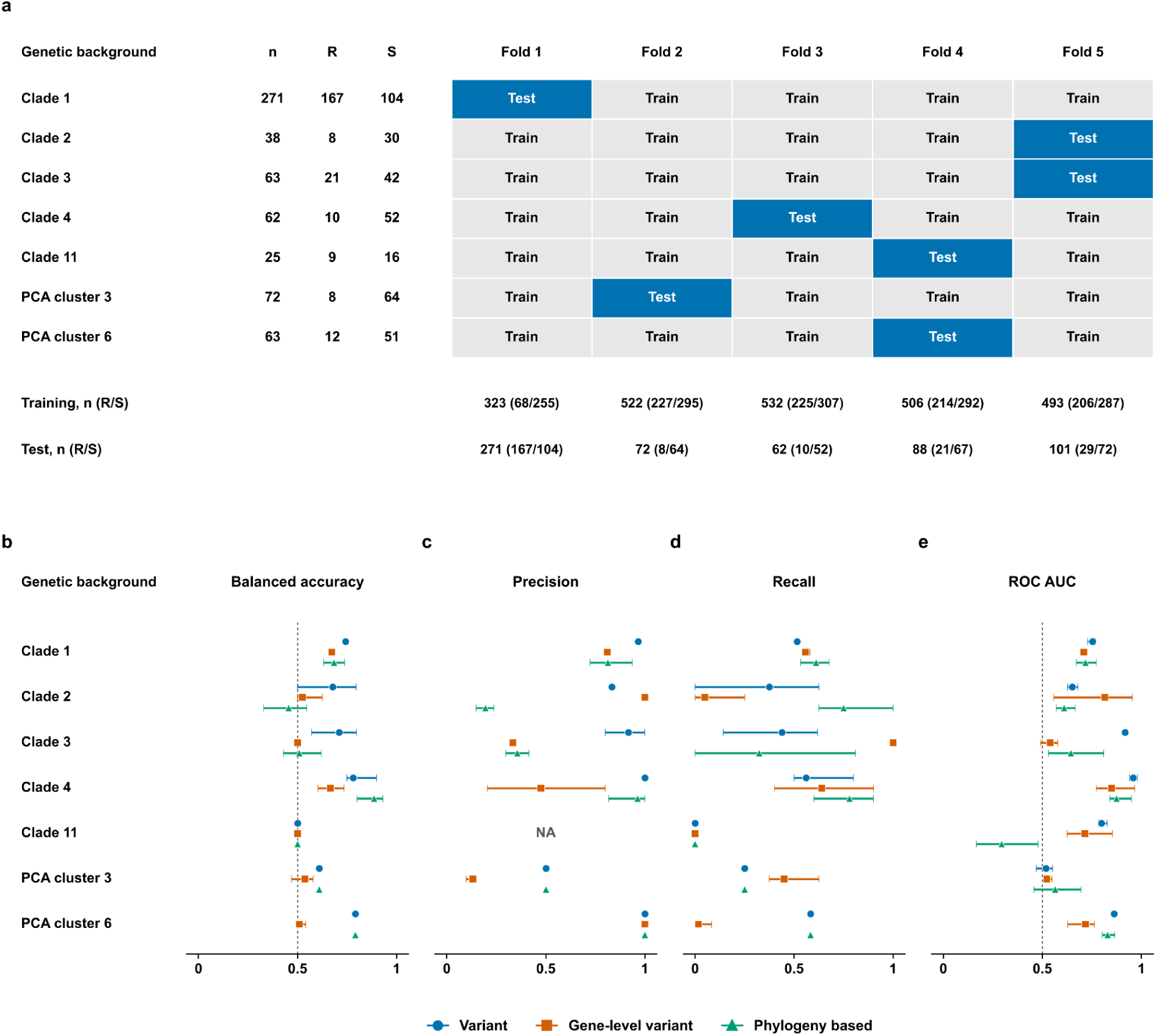
Fluconazole-resistance prediction across held-out genetic backgrounds. **a,** Training and test membership of seven genetic backgrounds in clade-grouped cross-validation. The cohort contains 594 isolates (235 resistant and 359 susceptible), from backgrounds with ≥15 isolates and ≥5 resistant isolates. Rows give total (n), resistant (R) and susceptible (S) isolate counts; columns show the five outer folds. Grey and blue cells indicate training and testing, respectively. Counts below the matrix give training and test totals, with R/S counts in parentheses. These assignments recur across all 5 repeats and are identical across feature sets. **b–e,** Balanced accuracy (**b**), precision (**c**), resistant-isolate recall (**d**) and ROC AUC (**e**), calculated separately within each held-out background. Blue circles, orange squares and green triangles denote variant, gene-level variant and phylogeny-based features, respectively. Points show means across 5 repeats; horizontal whiskers span the minimum and maximum. Classification thresholds were selected using training data. Balanced accuracy averages resistant- and susceptible-class recall; precision is the fraction of predicted-resistant isolates that are resistant; recall is the fraction of resistant isolates correctly identified. ROC AUC measures discrimination across prediction thresholds. Dashed lines at 0.5 mark chance-level balanced accuracy and ROC AUC. Precision is undefined when no isolate is predicted resistant; means omit undefined repeat values, and NA indicates no defined values in any repeat. Repeat-specific values and numbers of defined estimates are provided in the source data. Repeats reuse the same isolates and group assignments; whiskers describe variability across fit. Variant-based predictions outperformed phylogeny-based predictions in clades 2 and 3, whereas phylogeny-based predictions achieved the highest balanced accuracy and recall in clade 4. Both approaches performed similarly in PCA cluster 6. All three approaches classified every clade 11 isolate as susceptible, failing to detect resistance in this background. When clade 1 was withheld, variant-based predictions achieved the highest balanced accuracy and precision but detected fewer resistant isolates than the other approaches. No feature set therefore performed consistently well across all held-out backgrounds.

**Extended Data Figure 9.**
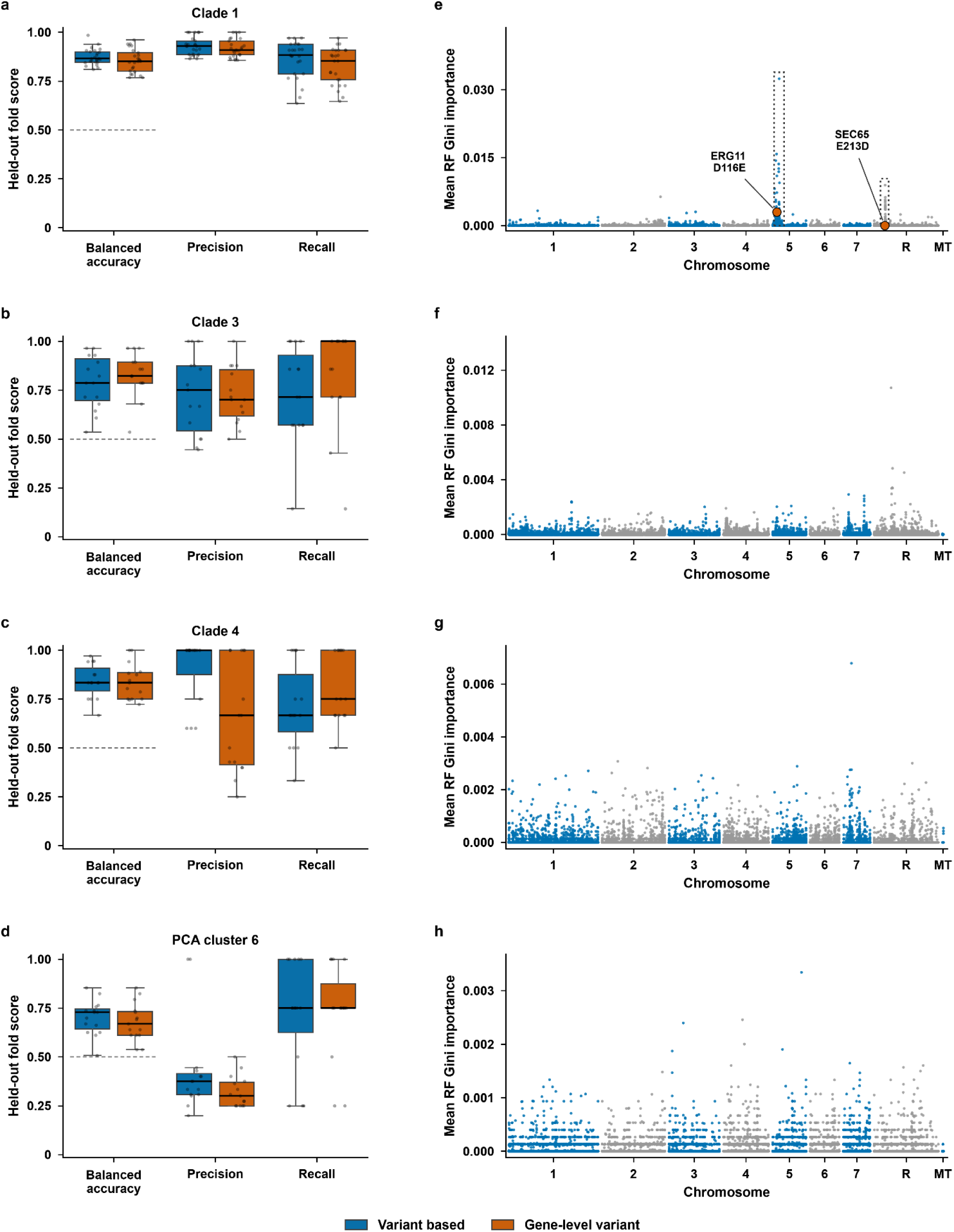
Within-group random-forest performance and variant-level feature importance for fluconazole-resistance prediction. Rows correspond to clade 1 (a, e; n = 271, including 167 resistant isolates), clade 3 (b, f; n = 63, 21 resistant), clade 4 (c, g; n = 62, 10 resistant) and PCA cluster 6 (d, h; n = 63, 12 resistant). **a–d,** Held-out balanced accuracy, precision and recall for variant-based (blue) and gene-level variant (orange) random forests trained separately within each genetic group. Violin widths show the distribution of fold estimates; internal lines mark the median and quartiles, and points represent individual held-out folds. Each input type was evaluated using five repeats of five-fold stratified cross-validation in clade 1 (25 estimates per violin) and five repeats of three-fold stratified cross-validation in the other groups (15 estimates per violin). Hyperparameters and classification thresholds were selected using inner cross-validation within each outer training set. Balanced accuracy is the mean of resistant- and susceptible-class recall; precision and recall refer to the resistant class. Short dashed lines beneath the balanced-accuracy violins mark 0.5. Fold distributions describe variability across repeated evaluations, not independent biological replicates. **e–h,** Mean normalized random-forest Gini importance of individual variant predictors, plotted by genomic position. Each point represents a variant retained in at least one outer-training model. Gini importance measures the normalized reduction in class impurity attributable to splits on a feature. Importance was averaged across all 25 or 15 outer-training models, assigning zero when a variant was not retained. Alternating blue and grey distinguish chromosomes; R denotes chromosome R and Mt denotes the mitochondrial genome. Genomic coordinates are aligned across groups, whereas importance-axis limits vary.

**Extended Data Figure 10.**
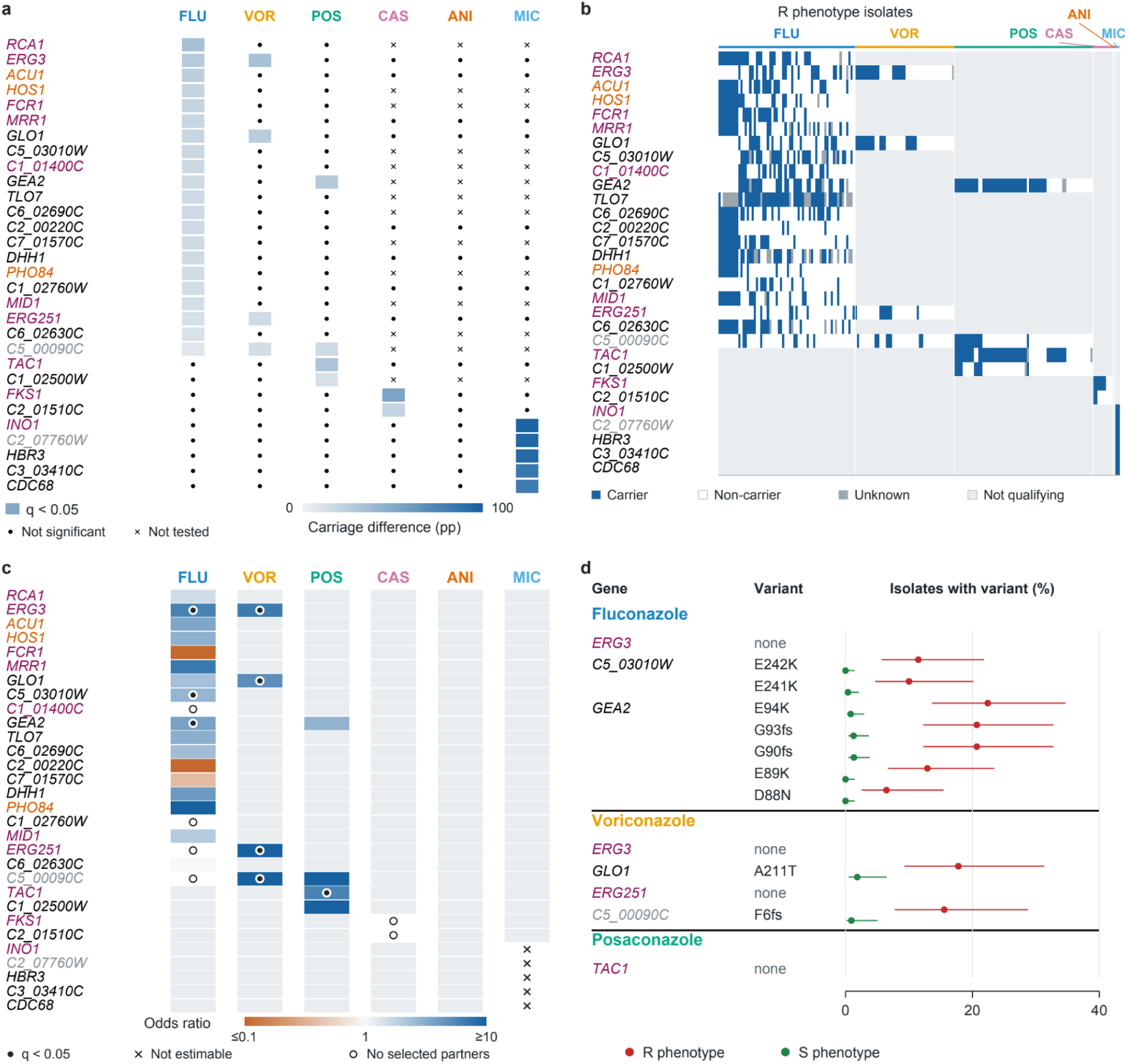
Prioritisation of resistance candidate mutations in isolates without high-confidence FungAMR substitutions. Cohort selection, variant eligibility and statistical analyses are described in Methods, “Prioritizing candidate genes.” Analyses included n = 471 distinct isolates. Eligible sample sizes (n, isolates; R phenotype/S phenotype) were: fluconazole (FLU), 62/268; voriconazole (VOR), 45/108; posaconazole (POS), 63/79; caspofungin (CAS), 9/374; anidulafungin (ANI), 1/192; and micafungin (MIC), 2/264. Isolates could contribute to multiple antifungal cohorts. R phenotype and S phenotype denote resistant and susceptible classifications, respectively; q denotes the Benjamini–Hochberg-adjusted P value. **a,** Initial gene enrichment. Rows show 30 genes and columns show six antifungals. Coloured cells represent 36 positively enriched associations (q <0.05); blue intensity shows the R phenotype minus S phenotype carrier percentage difference, in percentage points (pp). Black dots indicate comparisons that did not qualify, including non-estimable comparisons; crosses indicate combinations outside the original testing scope. **b,** Carriage of qualifying genes among R phenotype isolates. Rows match a; columns represent 182 isolate–antifungal observations from n = 92 distinct isolates, grouped by antifungal. Blue indicates carriage of an eligible variant; white, confirmed non-carriage; dark grey, unknown carriage because of missing calls; and pale grey, a gene–antifungal association that did not qualify in a. **c,** Gene enrichment after accounting for genetic background and selected co-carried genes, with axes matching a. Co-carriage denotes variants in different genes occurring in the same isolate. Colour shows the conditional odds ratio on a logarithmic scale: orange indicates enrichment in S phenotype, blue indicates enrichment in R phenotype, and near-white indicates equal odds. Colours saturate at odds ratios ≤0.1 and ≥10. Black dots indicate q <0.05; crosses, insufficient informative comparisons for estimation; open circles, no selected partner genes; and pale grey, associations outside the initial qualifying set. **d,** Individual-variant enrichment within the eight retained gene–antifungal associations. Rows show variants grouped by gene and antifungal; horizontal position shows the percentage of isolates with a known call carrying the variant. Red and green circles represent R phenotype and S phenotype, respectively, with 95% Wilson confidence intervals. All nine qualifying variants are shown. None indicates that no individual variant met the enrichment criteria. Protein changes use one-letter amino-acid codes; fs denotes a frameshift. Individual-variant tests accounted for genetic background; co-carriage adjustment applied to gene selection. Gene-name colours indicate prior evidence: magenta, published azole/echinocandin phenotype or drug-target evidence; orange, other-drug, cell-wall or indirect drug-response evidence; black, other informative annotated or predicted functions; and grey, uninformative annotations. Antifungal-name colours retain their original network assignments. Exact gene-, model- and variant-specific sample sizes and statistical results are provided in Source Data. Throughout, **q** is the BH-adjusted P value. All association tests were two-sided and exact, using Fisher’s exact test when only one informative stratum remained. Non-estimable tests contributed P = 1 solely when calculating multiple-testing corrections, while their reported P and q values remained missing. Gene-name colours retain the original networks’ evidence categories: magenta, published gene-specific azole/echinocandin phenotype (including tolerance) or drug-target evidence; orange, other-drug or cell-wall evidence, or qualified indirect drug responses; black, other informative annotated or predicted functions; grey, uninformative functional annotations. Antifungal-name colours identify the drugs and match their original networks. Exact gene- and model-specific sample sizes, carrier counts, effect estimates and confidence intervals are supplied in Source Data.

## Data and code sharing

Raw sequence reads of *C. albicans* and two *C. dubliniensis* isolates are available at NCBI SRA repository under BioProject PRJNA592373 and PRJNA1496987. Scripts used for the analyses are available at https://github.com/Landrylab/Jay_et_al_2026.

## Acknowledgements

We thank Jean-Barbeau and Annie Leduc for providing the University of Montreal Candida strain collection, and Alexandre K Dubé for helping with genomic DNA extractions. This project was funded by Genome Canada and Genome Québec grant (no. 6569) and by NSERC CREATE grant EvoFunPath to CRL and Genome Research and Development Initiative (GRDI) to AB. CRL holds the Canada Research Chair in Cellular Systems and Synthetic Biology.

## Authors contributions

Design research: CRL, AF, PD; Performed research: AJ, AP, CB, AF; Analysed the data: AJ, CB, AP, AF; Performed experiments: IGA, CB; Contributed DNA samples, phenotypic data and expertise: JK, LM, TD, SS, DGdL, AB, PD; Wrote the paper: AJ, CB, AF, IGA, CRL; Supervised the project: CRL, AF, IGA.

## Methods

### Library preparation and sequencing of isolates

Newly sequenced genomes in our study came from our lab at Université Laval, three public health labs across Canada (Alberta, Ontario and Québec) and from a personal collection at the Université de Montréal. Genomic DNA (gDNA) from clinical samples was extracted using either the Masterpure Yeast DNA Purification Kit (Lucigen) with specific modifications for yeasts (supplementary RNase treatment step) or following the ZymoBiomics DNA Miniprep kit instructions (ZymoResearch). gDNA was quantified with the Quant-iT™ PicoGreen® dsDNA Assay Kit (Life Technologies). Sequencing libraries were generated from 100 ng of gDNA using Illumina DNA Prep Kit (Illumina) and following manufacturer’s instructions. Library quantification was done with Kapa Library Quantification Kit - Complete Kit (Universal - Kapa Biosystems). Fragment mean size was determined using a Fragment Analyzer 5300 instrument (Agilent). Libraries were normalized and pooled, then denatured with 0.02 N NaOH and neutralized with a pre-load buffer. Pool was loaded at 170 pM on an Illumina NovaSeq X Plus 25B lane for 2x150 cycles (paired-end) following manufacturer’s instructions. A PhiX library was used as a control and mixed at 1% with gDNA libraries. Sample demultiplexing and FASTQ file generation were performed using the BCL Convert 4.2.4 program.

### Antifungal susceptibility classification and multidrug-resistance definitions

Antifungal susceptibility records from published studies and participating public health laboratories performed according to CLSI M27 or Yeast One (Y09; Remel/Oxoid) broth microdilution methodologies^83^ were compiled for fluconazole, voriconazole, posaconazole, caspofungin, micafungin and anidulafungin. Records were linked to isolate identifiers and represented using common drug-specific binary labels, encoded as R (1) and S (0). Across contributing sources, the final R labels corresponded exactly to MICs >2 µg ml⁻¹ for fluconazole, >0.125 µg ml⁻¹ for voriconazole, >0.06 µg ml⁻¹ for posaconazole and >0.25 µg ml⁻¹ for each of caspofungin, micafungin and anidulafungin. S labels corresponded to positive MICs at or below the respective thresholds. Quantitative MIC values were retained separately for the association analyses described below.

These binary labels represent an operational reduced-susceptibility endpoint. Relative to the *C. albicans* interpretive categories in CLSI M27M44S, fourth edition, the fluconazole threshold includes susceptible-dose-dependent values, while the voriconazole and echinocandin thresholds include intermediate values in the R category; these categories were not represented separately in the binary analyses^84^. The posaconazole threshold corresponds numerically to the *C. albicans* epidemiological cutoff in CLSI M57S, third edition^85^, which distinguishes wild-type from non-wild-type isolates^86^. R and S therefore denote the study’s binary phenotype categories and should be interpreted using the explicit thresholds above.

For the descriptive summaries, multidrug resistance (MDR) was defined operationally as an R phenotype for at least one azole and at least one echinocandin and was evaluated among the 809 isolates with results for at least one drug from each class. Within-class cross-resistance was defined as an R phenotype for at least two drugs in the same class, with denominators restricted to isolates tested against at least two azoles (n = 659) or at least two echinocandins (n = 632). Pan-azole and pan-echinocandin resistance required R phenotypes for all three drugs in the respective class and were evaluated only among isolates tested against all three azoles (n = 610) or all three echinocandins (n = 575). These categories described observed phenotype patterns without implying a shared resistance mechanism. Categories overlapped, and an isolate could contribute to more than one category. Percentages were calculated using the denominator specific to each category; missing results were not counted as the S phenotype (Extended Data Fig. 4).

### Genome processing

We collected paired-end Illumina whole-genome sequencing reads and corresponding MIC measurements for 1,201 isolates from seven published studies^8,16,18–22^. Isolate identifiers, genome accessions, MIC measurements and associated metadata are provided in Supplementary Table 1. Software was installed in Conda environments, primarily from the Bioconda repository^87^, and the complete workflow is available at https://doi.org/10.5281/zenodo.13741341. Raw reads were downloaded from the Sequence Read Archive using prefetch and fasterq-dump from SRA Toolkit v3.0.6^88^. Read lengths and quality-score encoding were inspected using SeqKit v2.5.1^89^. Adapters and low-quality bases were removed using Trimmomatic v0.39^89^ with ILLUMINACLIP:2:30:10 LEADING:3 TRAILING:3 SLIDINGWINDOW:4:15 MINLEN:31. Read quality was assessed before and after trimming using FastQC v0.12.1^90^ with reports summarized using MultiQC v1.15^91^. Trimmed reads were aligned to haplotype A of the *C. albicans* SC5314 Assembly 22 reference genome, release A22-s07-m01-r183, obtained from the *Candida* Genome Database^92^, using the mem algorithm in BWA-MEM2 v2.2.1^93^. Read-group information was assigned during alignment, with identifiers encoding the sequencing flowcell and lane and a consistent sample name assigned to runs belonging to the same isolate. Alignments were coordinate-sorted using SAMtools v1.17^94^, and samtools calmd-bAQr was used to recalculate MD and NM tags and apply base-alignment-quality adjustment. Duplicate reads were identified and removed using Picard MarkDuplicates v3.0.0^95^ with REMOVE_DUPLICATES=true. The resulting BAM files were indexed with SAMtools, and read-depth statistics were calculated using mosdepth v0.3.4^96^. Newly sequenced isolates were processed using the same workflow, except that reads were trimmed using fastp v0.24.0^96^ to remove adapters, low-quality bases, and poly-G and poly-X tails. The fastp options were -n 5 -q 15 -u 40 -l 31 --detect_adapter_for_pe -5 --cut_front_window_size 1 --cut_front_mean_quality 3 -r --cut_right_window_size 4 --cut_right_mean_quality 15 -g -x.

### Variant Calling and Joint Genotyping

Whole-genome variant discovery was performed using GATK v4.1.4.1^97,98^. For each isolate, HaplotypeCaller was run separately for each chromosome in GVCF mode with ploidy set to two and two native PairHMM threads. Chromosome-level gVCFs were combined into one gVCF per isolate using Picard GatherVcfs^95^. A sample map linking each isolate to its gVCF was used to import the cohort into a GenomicsDB datastore with GenomicsDBImport^99^. Joint genotyping was performed separately by chromosome using GenotypeGVCFs, producing chromosome-level multisample VCFs that were subsequently concatenated using BCFtools v1.15^94^. Reference-genome indexing and auxiliary file preparation were performed using SAMtools v1.21^94^. Computationally intensive steps were executed on a high-performance computing cluster managed with Slurm^100^. Chromosome-level jobs were parallelized using job arrays, with up to 18 CPU cores and 128 GB RAM allocated per task.

### Strain genome quality control and Variant filtration

Sample-level quality control and SNP filtering were performed using VCFtools v0.1.16^101^ and BCFtools/HTSlib v1.21^94^. Per-isolate genotype missingness was calculated using vcftools --missing-indv. Seven isolates with more than 10% missing genotype calls were excluded, six mislabelled samples were removed, and two *C. dubliniensis* isolates were excluded, reducing the cohort from 1,201 to 1,186 records. For isolates represented by multiple sequencing runs, the run with the highest mean read depth was retained and 33 redundant lower-coverage runs were removed, yielding a final cohort of 1,153 C. albicans isolates. The complete joint-call VCF was subset to these 1,153 isolates before SNP-specific site selection to construct a shared master variant catalogue for association and prediction analyses. Sequence SNP alleles and indel alleles with different reference and alternate lengths were retained, including eligible alleles at mixed SNP/indel and multiallelic sites. Symbolic alleles, the spanning-deletion allele `*`, reference-identical alleles and equal-length multinucleotide substitutions were excluded. Multinucleotide substitutions were not decomposed into constituent SNPs. No global allele-frequency, carrier-count, genotype-missingness, mapping-quality or VCF FILTER threshold was applied to the master catalogue, and eligible alternate alleles without carriers in the retained cohort were preserved.

Multiallelic records were represented as separate target-alternate-allele records. Alleles were normalized against the SC5314 Assembly 22 haplotype-A reference using BCFtools/HTSlib v1.24, including reference-allele validation, reference-based left alignment and minimal allele representation. Normalization realigned 34,514 allele records, with no reference mismatches or duplicate standardized alleles. Original multiallelic genotypes and their mappings to standardized alleles were retained. Complete diploid genotypes were encoded as zero, one or two copies of the specified alternate allele; other called alternate alleles contributed zero copies of that allele. Thus, zero dosage indicated absence of the target alternate allele and did not necessarily indicate an originally homozygous-reference genotype. Partially missing, fully missing and non-diploid calls were recorded as missing.

Site-level annotations, including AC, AN, AF, NS, F_MISSING and MAF, were recalculated across the 1,153-isolate cohort. The final joint VCF was compressed and indexed using bgzip and tabix^94^. It contained 2,016,116 alleles: 1,710,082 SNP alleles at 1,637,614 distinct normalized SNP positions and 306,034 indel alleles at 235,477 distinct normalized indel positions. These comprised 1,707,132 nuclear and 2,950 mitochondrial SNP alleles, and 305,306 nuclear and 728 mitochondrial indel alleles. Among the indels, 188,113 had a longer alternate allele and 117,921 had a shorter alternate allele. Subsequent sample selection and variant filtering were applied according to the analysis-specific rules described below.

### Phylogenetic Tree Building

To avoid representing decomposed multiallelic sites as multiple alignment columns, phylogenetic inference used a separate, undecomposed, one-record-per-position biallelic SNP panel. The resulting PHYLIP alignment contained 1,153 samples and 957,775 distinct genomic positions. Heterozygous genotypes were represented by IUPAC ambiguity codes and missing genotypes by N. Maximum-likelihood inference was performed with IQ-TREE v2.1.3^102,103^ under the GTR+F+I+R6 substitution model. GTR allows the six nucleotide-substitution rates to differ, +F uses empirical nucleotide frequencies, +I estimates a proportion of invariant sites and +R6 models among-site rate heterogeneity using six FreeRate categories rather than a discrete gamma distribution^104^. This model was specified because the alignment exhibited both substantial compositional and rate heterogeneity and contained 504,394 constant positions (52.7% of the alignment), making explicit modelling of invariant and rate-variable sites appropriate. The maximum-likelihood search was run with eight CPU threads and random seed 20250812. Two *Candida dubliniensis* genomes were specified as outgroups. Branch support on the resulting maximum-likelihood topology was evaluated using 1,000 replicates of the Shimodaira–Hasegawa-like approximate likelihood-ratio test (SH-aLRT)^105^. Phylogenetic trees were visualized and annotated using Interactive Tree of Life v6 (iTOL)^106^.

### Linkage disequilibrium (LD) pruning of the SNP dataset

Nuclear SNPs from the master variant catalogue for the 1,153 *C. albicans* isolates were restricted to a minor allele frequency of at least 0.05 and genotype missingness of no more than 10%, retaining 243,594 SNPs. Mitochondrial SNPs were excluded. Genotypes were stored in PLINK binary format, with variant identifiers specifying chromosome, position, reference allele and alternate allele. Linkage disequilibrium was estimated using the squared genotype-correlation coefficient, (r^2^)^107^, and SNPs were pruned using PLINK v1.9.0-b.7.11^108^ with --indep-pairwise 50 5 0.2. This procedure considered windows of 50 variants, advanced the window by five variants and iteratively removed variants involved in pairs with (r^2^ > 0.2). LD pruning removed 220,685 SNPs and retained 22,909 SNPs, which constituted the final LD-pruned SNP panel.

### Principal Component Analysis (PCA) of SNPs

Using the LD-pruned SNP set diploid genotypes were encoded additively as the number of alternate-allele copies: 0 for homozygous reference, 1 for heterozygous and 2 for homozygous alternate genotypes. PCA was performed directly in PLINK v1.90b5.3 using the --pca 50 option^108,109^. PLINK mean-centered the genotype dosages and scaled each SNP according to its allele-frequency variance to construct a variance-standardized genomic relationship matrix, from which the first 50 PCs were extracted. The first 20 PC scores were centered and scaled in R v4.2.1^110^ before clustering. PCA and clustering figures were generated with ggplot2 v3.4.0^111^.

### Unsupervised clustering

To obtain an unsupervised genetic clustering of the samples, PC1–PC20 were centred and scaled before k-means clustering using the Hartigan–Wong algorithm^112^ across K=2–30 (seed 42; 150 random starts). The number of clusters was evaluated using mean silhouette width^113^ and the gap statistic with 150 reference simulations and the firstSEmax rule^114^. Silhouette width measures whether samples are closer to members of their own cluster than to members of neighbouring clusters, with higher values indicating clearer separation. The gap statistic compares the observed clustering with that obtained from simulated data lacking clear cluster structure, and the firstSEmax rule selects the smallest K that performs close to the best solution. The maximum silhouette width occurred at K=23, whereas the gap statistic selected K=2. These candidate solutions were compared using the Calinski–Harabasz^115^ and Davies–Bouldin indices^116^. The Calinski–Harabasz index compares the separation between clusters with the spread of samples within clusters, with higher values indicating a better solution. The Davies–Bouldin index measures how similar each cluster is to its nearest neighbouring cluster, with lower values indicating better separation. Stability was also assessed using the mean adjusted Rand index (ARI)^117^ across 20 independent 80% row subsamples drawn without replacement. The ARI measures how consistently two analyses group the same samples together after accounting for chance, with 1 indicating identical groupings and values near 0 indicating chance-level agreement. Clusterwise Jaccard stability^118^ was calculated over 150 bootstrap replicates. This score measures how consistently the members of each cluster remain together across repeated analyses, with values approaching 1 indicating a highly stable cluster. The effect of PC selection was assessed by repeating the clustering with 200 random subsets containing 80% of the PCs and comparing each result with the full-PC solution using the ARI. A prespecified rule selected the higher K when cluster separation improved while the row-subsample ARI, mean Jaccard stability and PC-subsampling ARI were each retained within 0.02 of the lower-K solution. This procedure selected K=23 as the final consensus. Final cluster assignments were obtained from a K=23 k-means fit using seed 42 and 150 random starts, and final clusterwise stability was evaluated over 200 bootstrap replicates.

### ADMIXTURE analysis

Model-based ancestry proportions were estimated from the LD-pruned SNPs using ADMIXTURE v1.3.0^119,120^. Models were fitted for K=2–30, where K represents the assumed number of ancestral populations. Ten independent runs were performed for each K using recorded random seeds and tenfold cross-validation. Cross-validation withholds part of the genotype data and measures how accurately the fitted model predicts it, with lower error indicating better fit. For each K, the run with the lowest cross-validation error was used to display ancestry proportions, while results from all ten runs were summarized to assess consistency.

### Clade annotation

Published clade names followed the established *C. albicans* MLST nomenclature^121^. The published clade labels available for 139 isolates were summarized within each K=23 PCA cluster. ADMIXTURE profiles were considered together with phylogenetic position and K=23 PCA-cluster membership when extending published clade labels to previously unlabelled isolates. A fixed cluster-to-clade map was then used to extend a named clade to unlabelled members of clusters with a supported published-clade match, with phylogenetic placement and ADMIXTURE ancestry profiles used as supporting checks. Clusters without a supported named-clade assignment were retained as “Other”. The “NC” label for the new clade described by Sitterle et al.^16^ was retained when displaying the literature-labelled subset but was not extended to unlabelled isolates.

### LOH detection and quantification

At each genomic position, a genotype was considered callable when both alleles were present and heterozygous when the two called alleles differed; records sharing the same genomic position were counted once. For each isolate, callable and heterozygous SNP positions were counted in consecutive, non-overlapping 10-kb windows across the eight nuclear chromosomes; mitochondrial variants were excluded. This SNP-based approach follows previous genome-wide analyses of LOH in *C. albicans^19,122^*.

A window was classified as having low heterozygosity when it contained at least ten callable SNP positions and no more than 10 heterozygous positions. Candidate LOH tracts contained at least three low-heterozygosity windows, allowed no more than three consecutive intervening windows that did not meet this definition and required at least 80% of the windows in the final tract to have low heterozygosity. Non-low terminal windows were removed when necessary to meet this purity threshold, and tracts were not allowed to span chromosomes. LOH coverage per isolate was calculated by merging overlapping tracts, clipping coverage to the chromosome lengths and dividing the covered length by the 14,282,759-bp reference genome. Median copy number and baseline ploidy were recorded from overlapping 10-kb copy-number bins for annotation but were not used to call LOH.

To identify phylogenetically shared candidate tracts, each tract was compared with tracts in the two neighbouring samples on either side of the same isolate in the ladderized tree display order. A neighbouring tract was considered a match when it occurred on the same chromosome and had an interval Jaccard similarity — intersection length divided by union length — of at least 0.20^123^. A tract was classified as candidate synapomorphic when at least 80% of the available neighbouring samples carried a matching tract. “Candidate” is used because this display-neighbourhood rule identifies phylogenetically shared segments but does not constitute formal ancestral-state reconstruction.

For clade-level characterization, the analysis was restricted to the 861 isolates with clade annotations. Within each clade and chromosome, candidate tracts were grouped into connected regions when they overlapped or were separated by no more than 10 kb, their midpoints differed by no more than 150 kb and their lengths differed by no more than threefold. Consensus starts and ends were defined as the 25th percentile of tract starts and the 75th percentile of tract ends, respectively. Regions sharing an identical consensus start or end were reduced to the region with the greatest clade-support fraction, followed by the number of supporting isolates and region length. Regions were flagged for reporting when they occurred in more than 30% of the isolates in a clade and were longer than 100 kb.

### Whole-genome ploidy and CNV detection

To detect whole-genome ploidy changes, whole-chromosome aneuploidies and gene-overlapping copy-number variation, GC-content bias was corrected against the SC5314 reference genome using computeGCBias and correctGCBias from deepTools v3.5.1^124^, which implement the fragment-level correction described by Benjamini and Speed et. al. 2012^125^. Corrected alignments were summarized in non-overlapping 1-kb bins using SAMtools v1.15^126^ and BEDTools v2.30.0^127^. Bins were excluded when fewer than 75% of bases were covered, median matched-base depth was below 80, or the bin overlapped rDNA, a centromere, a telomere or an annotated repeat. Mitochondrial bins were excluded. The remaining chromosome-end trend was corrected by modelling log-transformed depth as a function of distance to the nearest chromosome end using quadratic locally weighted regression (LOESS; span 0.3)^128^. Expected depth was obtained by tenfold cross-validation, and each observed value was divided by its held-out prediction and rescaled to the original genome-wide median.

Copy-number calling combined corrected read depth with allele fractions at heterozygous biallelic SNPs, following the joint depth-and-allele-balance framework used for *Candida* genome analysis in YMAP^129^. For each isolate, depth was expressed as the log2 ratio relative to the sample-wide median, and allele fraction as the absolute deviation from 0.5. Circular binary segmentation was applied separately to the two tracks with DNAcopy (alpha = 0.01; minimum width = three observations)^130,131^. Breakpoints from both segmentations were combined to define consensus intervals. Each interval was summarized by its coordinates, median log2 depth ratio, number of informative SNPs, median allele fraction and nearest expected allele-fraction state. Allele-fraction evidence was included in copy-number scoring when at least ten informative SNPs were present.

Baseline ploidy was inferred from genome-wide allele-fraction peaks together with the depth behaviour of copy-neutral segments. Integer segment copy number was assigned by jointly comparing the observed depth ratio and allele fraction with those expected under candidate copy-number states. Chromosome copy number was the length-weighted modal segment state, and a gene was assigned the copy number of a segment covering at least 50% of its length. Ambiguous automated profiles were reviewed using the depth, allele-fraction and breakpoint plots. The final manually reviewed calls reclassified two isolates from 2N to 4N. The resulting 1,153-isolate collection comprised 1,127 2N (97.7%), eight 3N (0.7%), 11 4N (1.0%) and seven 6N (0.6%) isolates.

For the genome-wide display, 1-kb calls were grouped into non-overlapping 10-kb windows. A gain or loss was retained when at least 70% of available 1-kb bins supported the same direction relative to the isolate baseline; otherwise, the window was assigned the baseline copy number. The displayed state was the rounded median absolute copy number among supporting bins. Whole-chromosome events were defined by a chromosome copy number different from the isolate baseline. Among the 1,127 diploid isolates, segmental events were defined as consecutive altered gene-overlapping loci spanning at least 30 kb on a chromosome without a whole-chromosome event in the same isolate.

### Validation of selected polyploids

Selected *C. albicans* isolates were assessed by flow cytometric measurement of relative cellular DNA content to evaluate genome-inferred ploidy. Comparison with reference strains of known ploidy provides an established approach for assessing fungal DNA content^132^. The assay panel comprised 23 isolates from our study and two diploid *C. albicans* controls, fRS1 and fRS757. Haploid, diploid, triploid and tetraploid *Saccharomyces cerevisiae* reference strains were included as supplementary controls. DNA content for all isolates and controls was stained with SYTOX™ green (Thermo Fisher, Waltham, USA) according to the procedure described in Bautista et al. (2024)^81^.

Fluorescence was acquired using a Guava easyCyte HT flow cytometer with InCyte software version 3.4, targeting 5,000 events per well. Linear green-fluorescence pulse-height distributions (GRN-B-HLin) were visualized as a heatmap showing the percentage of all recorded events in each 500-unit fluorescence bin, using a common log-transformed colour scale (Extended Data Fig. 3e). Ploidy was assessed by visual comparison of the fluorescence profiles with the diploid *C. albicans* reference fRS1 and with the additional reference strains providing supporting comparisons. The tested panel included the samples BL014, BL042 and BL205, corresponding to the diploid, triploid and tetraploid genomic examples shown in Extended Data Fig. 3a–c.

### FungAMR annotation and antifungal susceptibility

Protein-changing alleles in the standardized variant catalogue were matched to *C. albicans* records in the archived FungAMR database using the reference gene and exact amino-acid substitution. Gene names and aliases were harmonized to reference identifiers before matching, including CYP51/CYP51A to *ERG11* and FKS/GSC1 to *FKS1*. All allele-specific transcript annotations were considered. For each gene–substitution–antifungal combination, resistance and sensitivity evidence was resolved jointly before applying confidence thresholds. Positive scores denote resistance evidence and negative scores denote sensitivity evidence; smaller absolute values indicate greater confidence. We retained the nonzero score with the smallest absolute value, preferring the positive resistance score when absolute values were equal. Thus, scores +8 and −2 resolve to −2, whereas +2 and −2 resolve to +2. High-confidence resistance substitutions were defined by resolved scores of +1 to +4 for the corresponding drug. The Erg11 V488I and D116E substitutions were excluded from matching due to it being a known error in the database.

For the evidence-coverage analysis, observed substitutions in Erg11, Upc2, Cdr1, Cdr2, Mdr1, Tac1, Mrr1, Mrr2, Cap1 and Fks1 were compared with all available FungAMR drug annotations. Evidence for an exact single substitution was resolved separately from evidence for a multiple-substitution combination. Membership in a combination of mutations that had a FungAMR score did not constitute independent evidence for the effect of an individual mutation in that combination so a combination score did not replace a component’s standalone score, even when the combination had higher-confidence evidence. After resolving conflicting scores within each drug and matching context, each observed substitution was assigned once to the first applicable category: high-confidence resistance evidence (+1 to +4), lower-confidence resistance evidence (+5 to +8), membership in a resistance-associated multiple-substitution record, sensitivity-associated evidence (-1 to -8), or no matching record. This coverage summary combined evidence across drugs; phenotypic concordance analyses retained drug-specific annotations and used only exact single-substitution matches.

An isolate was classified as a carrier when its called genotype contained at least one copy of the alternate allele encoding a qualifying substitution. Duplicate annotations and database records were consolidated, and each isolate was counted once per substitution and once per drug-specific carrier group, irrespective of the number of qualifying alleles. One and two copies of the specified alternate allele in diploid genotype calls defined heterozygous and homozygous-alternate states, respectively. Gene-overlapping LOH was recorded when a called LOH tract overlapped the corresponding reference gene interval by at least one base. This definition did not require the entire gene or the substitution itself to lie within the tract (Methods, LOH detection and quantification). All matched alleles used in the phenotype assessment were callable in all 1,153 isolates.

For phenotypic concordance, we excluded the 370 isolates from Gong et al. (2023)^20^ and 14 from Chew et al. (2021)^18^, because these studies contributed isolates to the FungAMR evidence base. Of the 769 remaining isolates, 494 had at least one binary susceptibility phenotype. Analyses included only isolates with an available resistant or susceptible classification for the corresponding drug: 487 for fluconazole, 289 for voriconazole, 241 for posaconazole, 396 for caspofungin, 205 for anidulafungin and 283 for micafungin. A drug-specific carrier required at least one exact single-substitution match with a resolved score of +1 or +2 for that drug. Carrier and non-carrier percentages used their respective group totals as denominators. Gene-contribution bars used all resistant isolates tested against the corresponding drug as the denominator; isolates could contribute to more than one gene.

Associations between substitution carriage and resistance were evaluated using two-sided Fisher’s exact tests^24^ on 2 × 2 tables with carrier/non-carrier rows and resistant/susceptible columns. Sample odds ratios were calculated from these counts. *P* values were adjusted using the Benjamini–Hochberg procedure^25^ across the five drugs with nonzero row and column margins and are reported as *q* values. Anidulafungin was not testable because no eligible isolate carried a drug-matched high-confidence resistance substitution. The resolved evidence and phenotype-assessment results are provided in Supplementary Table 3.

### Copy-number variation and antifungal susceptibility

Gene-level copy-number variation (CNV) was compared with antifungal susceptibility among all 1,127 isolates assigned a diploid baseline ploidy (Methods, Whole-genome ploidy and CNV detection). Analyses included *ERG11*, *TAC1*, *MRR1*, *MRR2*, *UPC2*, *MDR1*, *CDR1*, *CDR2*, *CAP1* and *FKS1* genes. High-confidence resistance substitutions were defined as exact single-substitution matches with resolved FungAMR scores of +1 or +2, with evidence resolution and alternate-allele carriage determined as described above (Methods, FungAMR annotation and antifungal susceptibility).

For comparisons between copy-number gain carriers and non-carriers (Extended Data Fig. 6d,e), we excluded 206 diploid isolates carrying at least one high-confidence substitution in any gene for any drug represented in FungAMR, leaving 921 eligible isolates before restriction by phenotype availability. For each gene, gain carriers had an absolute copy number greater than two, and non-carriers comprised eligible isolates without a called CNV at that gene. Isolates with copy-number losses at the focal gene were excluded from these comparisons. Each isolate was counted once per gene and comparison group. Analyses were conducted separately for fluconazole, voriconazole, posaconazole, caspofungin, micafungin and anidulafungin. Resistance fractions used isolates with an available binary resistant or susceptible classification, whereas MIC summaries used isolates with an available numerical log₂ MIC for the corresponding drug. Resistance differences were calculated as 100 times the gain-carrier resistance fraction minus the non-carrier resistance fraction and expressed in percentage points. MIC differences were calculated as the median log₂ MIC among gain carriers minus that among non-carriers. Comparisons were displayed when at least two gain carriers and at least one non-carrier had the relevant phenotype measurement.

To examine CNV effects within established mutation backgrounds (Extended Data Fig. 6f), we selected diploid carriers of each *ERG11* substitution with a resolved high-confidence resistance annotation for the corresponding drug. Isolates carrying a high-confidence substitution in another gene for any drug were excluded; additional *ERG11* substitutions were permitted. Eligible isolates were divided into carriers with an *ERG11* CNV and carriers of the same substitution without an *ERG11* CNV. The difference in median log₂ MIC was calculated as the median in the CNV-plus-substitution group minus the median in the substitution-only group. Comparisons were displayed when both groups contained at least two isolates with available MICs. Isolates carrying multiple qualifying substitutions could contribute to more than one comparison. Group sizes, resistance counts, median MICs and observed differences are provided in Supplementary Table 2 and the source data for Extended Data Fig. 6.

### Genome-wide Association Study of SNPs and Indels

We tested nuclear SNPs and indels for association with antifungal minimum inhibitory concentration (MIC), using the retained isolates and joint variant calls described above. Analyses used the recorded log₂ MIC values as quantitative phenotypes, such that a one-unit increase corresponds to a doubling of MIC. Isolates with missing values were excluded separately for each drug, yielding 871 isolates for fluconazole, 659 for voriconazole, 611 for posaconazole, 766 for caspofungin, 667 for micafungin and 589 for anidulafungin.

Genotypes were stored in PLINK binary format using PLINK v1.9.0-b.8^108^ and analysed as additive dosages of zero, one or two copies of the tested allele. The effect allele was recorded for each association. Variant eligibility was determined within each drug-specific cohort. Variants required a minor allele frequency (MAF) of at least 0.005, calculated from called alleles, genotype missingness of no more than 10%. After filtering, the numbers of tested SNPs and indels combined were 463,959 for fluconazole, 464,880 for voriconazole, 461,365 for posaconazole, 467,792 for caspofungin, 462,935 for micafungin and 467,259 for anidulafungin.

Associations were fitted using the mixed linear model implemented in GCTA v1.95.0^134,135^. Each model included an intercept, the additive effect of the tested variant and a random genetic effect accounting for relatedness among isolates. Relatedness was represented by a genetic relationship matrix constructed from 20,993 common nuclear SNPs selected using the LD-pruning parameters described above and retained at MAF ≥0.05 and genotype missingness ≤10% across all 1,153 isolates. Genotype dosages were centred using full-cohort allele frequencies and divided by their expected standard deviation, √[2p(1−p)], where p is the frequency of the coded allele. Missing dosages were replaced by their mean, and the matrix was calculated as the standardized-genotype cross-product divided by the number of retained SNPs. The matrix included all eight nuclear chromosomes and was subset to each drug cohort. Genetic and residual variances were estimated under the null model by restricted maximum likelihood and held fixed during variant testing. Missing genotypes at tested variants were replaced by their drug-cohort mean dosage.

For each variant, the null hypothesis of no allelic effect was tested using the squared effect estimate divided by its squared standard error, referenced to a chi-squared distribution with one degree of freedom. The resulting two-sided P value is the probability, assuming no variant effect and the fitted model’s assumptions, of obtaining a test statistic at least as extreme as the observed statistic. Genome-wide significance was defined separately for each drug as P < 0.05/M, where M is the total number of tested SNP and indel alleles. This Bonferroni threshold controls the probability of at least one false-positive result within a drug scan at 5%, provided the individual null P values are valid. A suggestive threshold of P < 1/M corresponds to one expected false-positive test per scan under a calibrated global null. Suggestive associations were treated as candidates rather than genome-wide significant findings.

Manhattan plots displayed −log₁₀(P) against genomic position, with the two drug-specific thresholds indicated. Quantile–quantile (Q–Q) plots compared ordered observed −log₁₀(P) values with those expected under a uniform null P-value distribution, using expected probabilities of (r−0.5)/M for ranks r = 1,…,M. The diagonal indicates agreement between observed and expected values. The genomic inflation factor, λGC, was calculated as the median one-degree-of-freedom chi-squared statistic derived from the association P values divided by its theoretical null median, approximately 0.454936^136^. Values near one indicate agreement at the median; values above or below one indicate inflation or deflation, respectively. All tested variants were included. These diagnostics describe the test-statistic distribution but do not establish calibration of its most extreme tail. P values were not rescaled using λGC.

### Gene-level variant GWAS

We combined rare variants within genes to test whether carriage of potentially functional variation was associated with antifungal MIC^37^. The SNP/indel genotype panels described for the variant GWAS were used before application of its association-specific frequency and carrier-support filters. Allele-specific consequences were obtained using SnpEff v5.2^137^ with the SC5314 reference annotation, retaining variants classified as HIGH or MODERATE impact. These categories include predicted disruptive changes, such as frameshifts and premature stop codons, and changes such as amino-acid substitutions and in-frame indels. Each allele was assigned one matching nuclear-gene annotation, prioritizing HIGH over MODERATE impact and then annotations with a protein-level consequence; remaining ties were resolved deterministically by gene identifier and consequence. Alternate alleles were matched to their own annotations, using exact sequence matching or an unambiguous trimmed indel representation.

Variant selection was repeated within each of the six drug-specific cohorts defined above. A variant qualified when it was carried by at least one isolate and by no more than 5% of the total cohort. This carrier frequency counts isolates with at least one copy of the alternate allele, rather than the proportion of called chromosomes carrying that allele. Qualifying variants were collapsed into a binary gene-level indicator: an isolate was a carrier if it carried any qualifying variant, regardless of zygosity or the number of variants present. It was a noncarrier only when all qualifying variants were called and absent. If none was detected but at least one constituent genotype was missing, the gene-level indicator was missing. An observed qualifying variant therefore took precedence over missing calls at other sites.

Gene markers were tested when both carriers and called noncarriers were present. No lower variant-frequency or gene-marker-frequency threshold, additional minimum carrier count, or additional genotype-missingness threshold was imposed. The numbers of tested genes were 6,294 for fluconazole, 6,282 for voriconazole, 6,281 for posaconazole, 6,290 for caspofungin, 6,285 for micafungin and 6,281 for anidulafungin. For GCTA input, noncarriers and carriers were represented by dosages of zero and two, respectively; missing indicators remained missing. Reported effect estimates and standard errors were multiplied by two to express the carrier–noncarrier difference in log₂ MIC. This rescaling does not change the association P value.

Associations used a leave-one-chromosome-out (LOCO) relatedness adjustment^135^. Starting from the same common-SNP panel and full-cohort genotype standardization described for the variant GWAS, we constructed a separate relationship matrix for each nuclear chromosome by excluding SNPs on that chromosome and dividing the remaining genotype cross-product by the number of retained SNPs. Each gene was tested with the matrix excluding its own chromosome, reducing the contribution of nearby linked variants to the relatedness adjustment. Null-model variance components were estimated separately for each drug and excluded chromosome. Phenotype handling, other mixed-model settings, association *P*-value calculation, and Manhattan and Q–Q plotting followed the variant GWAS procedure. Bonferroni and exploratory thresholds were calculated as 0.05/M and 1/M, respectively, with M now denoting the number of tested genes for that drug. λGC was calculated as previously defined, using the gene-level squared effect-to-standard-error ratios as the observed chi-squared statistics.

### Network of leading gene-level variant associations

For each antifungal, genes were ranked by increasing gene-level association P value, with ties resolved by gene identifier. The 20 highest-ranked genes were retained irrespective of whether they crossed the genome-wide significance or suggestive thresholds. Combining the six lists produced 120 drug–gene memberships involving 91 distinct loci. Each connection linked a gene to a drug for which it appeared in the top-20 list. Connection colours identified fluconazole (blue), voriconazole (gold), posaconazole (green), caspofungin (pink), micafungin (light blue) and anidulafungin (orange). Gene-marker area was proportional to the number of drug-specific lists containing that gene. Genes represented in both azole and echinocandin lists were displayed centrally with outlined markers. These connections summarize ranked associations and do not represent measured molecular interactions.

Gene names, systematic identifiers, aliases and reference functional descriptions were obtained from the Candida Genome Database annotation for SC5314 Assembly 22, release A22-s07-m01-r183^92^. Descriptions were extracted from the gene-feature Note attributes in the reference GFF file and reviewed alongside gene-specific primary literature. Systematic identifiers and ORF aliases were used to distinguish similarly named loci and paralogues. Functions inferred from orthology or sequence features were described as predicted or putative. Drug-response annotations were checked against the species, agent and experimental context reported in the source study; original supplementary datasets from chemogenomic and transcriptional-regulator screens were consulted where required^41,138^.

Gene-marker and annotation colours summarized the type of prior evidence (Fig. 3c). Magenta indicated published azole- or echinocandin-response phenotypes or established antifungal-target evidence. Orange indicated evidence involving other agents, cell-wall phenotypes or indirect drug responses. Black indicated other annotated or predicted functions, and grey indicated that the reference description lacked an informative functional assignment. These categories were assigned independently of association rank or statistical significance. They describe prior biological evidence for a gene and do not establish a causal role for the specific variants aggregated in this study. The same magenta and orange categories identified literature-supported genes among the drug-specific top-20 associations labelled in the gene-level Manhattan plots.

### Random-forest prediction of fluconazole resistance

We compared three genomic input representations for predicting binary fluconazole resistance in the 871 phenotyped isolates described above, encoding resistant and susceptible isolates as 1 and 0, respectively. The input representations were prepared as follows.

#### Variant-based input

SNP genotypes were encoded as zero, one or two copies of the allele designated A2 in the PLINK record, with missing calls retained for subsequent imputation.

#### Gene-level variant input

Allele-specific functional annotation, the selection of HIGH/MODERATE-impact variants carried by no more than 5% of isolates, and aggregation into gene-level carrier indicators followed the gene-level variant GWAS procedure. Here, the carrier-frequency threshold was applied in the full binary-phenotyped fluconazole cohort before cross-validation. Indicators were encoded as 1 for carriage of at least one qualifying variant and 0 for called absence; missing indicators followed the previously described rule. These gene definitions were held fixed across validation folds.

#### Phylogeny-based input

Population structure was represented by SNP-derived PCs from the LD-pruned SNP set described previously. Within each training partition, missing dosages were replaced by the training-marker mean, markers were centred and scaled to unit variance, and PCA was fitted using randomized singular-value decomposition. Validation and test isolates were projected using the transformations fitted to their corresponding training samples. The number of retained PCs was selected from 2, 5, 10, 20, 40 and 80. Thus, the phylogeny-based representation summarized genome-wide relatedness through PCs rather than using tree branches as predictors.

All three representations were analysed using random-forest classifiers^139^ implemented in Python with scikit-learn^140^. Each forest comprised 500 decision trees trained on bootstrap samples, with random subsets of features considered at each split and Gini impurity used as the splitting criterion. Class weights were inversely proportional to class frequency within each tree’s bootstrap sample (balanced_subsample). The predicted resistance probability was the mean of the trees’ class probabilities.

Generalization was evaluated using five repeats of five-fold outer cross-validation. Stratified validation approximately preserved the resistant/susceptible ratio in each fold. Clade-grouped validation used the previously defined K=23 PCA clusters, keeping all members of a cluster together while approximately balancing classes across folds. Consequently, every held-out cluster was absent from its corresponding training set. The same outer partitions were used for all three input representations. Stratified folds used random seed 42; grouped repeats used seeds 42–46.

For variant and gene inputs, feature screening was performed separately in each cross-validation split using only the training samples, without access to the held-out test samples. Features with more than 10% missing calls or no variation among called genotypes were excluded, with no additional lower frequency threshold. Missing values were temporarily replaced by their training-set modal value to calculate scikit-learn’s chi-squared feature score against the binary phenotype. Features were ranked by decreasing score, with ties favouring higher training frequency, and up to 10,000 were retained. This selected feature set was held fixed during inner tuning. Model imputation was fitted separately within each inner training partition and again on the complete outer training set, using the most frequent value. For PC models, imputation, scaling and PCA were refitted within every inner training partition and on the complete outer training set.

Hyperparameters were selected by three-fold inner cross-validation using the same stratified or cluster-grouped design as the outer analysis. Variant and gene models evaluated nine ordered combinations of maximum features considered per split, minimum samples per leaf and maximum tree depth: (√p, 2, unlimited), (√p, 1, unlimited), (√p, 5, unlimited), (√p, 10, unlimited), (√p, 2, 20), (0.05p, 2, unlimited), (0.10p, 2, unlimited), (0.20p, 2, unlimited) and (0.10p, 5, 20), where p is the number of input features. PC models crossed the six candidate PC counts with the first, third and eighth forest configurations. Within each inner partition, 80 PCs were fitted once and subset for these candidates; the final outer-training PCA fit used the selected component count. The configuration with the highest mean inner-fold receiver operating characteristic area under the curve (ROC AUC) was selected; ties were resolved by the stated candidate order. Inner splits and forests used seed 42 plus the outer-fold number.

A classification threshold was chosen from the selected configuration’s pooled inner out-of-fold probabilities to maximize balanced accuracy. Candidate thresholds comprised the observed probabilities and 0, 0.5 and 1; ties favoured the value closest to 0.5 and then the lower value. The selected model was refitted on all outer training samples and evaluated once on the held-out fold, classifying probabilities at or above the threshold as resistant. ROC AUC measured discrimination across thresholds; balanced accuracy was the mean of resistant- and susceptible-class recall. Precision was the proportion of predicted-resistant isolates that were resistant, and recall was the proportion of resistant isolates correctly identified. At fold level, undefined precision or recall was recorded as zero.

Performance was summarized across the 25 outer folds per representation and validation design. Mean ROC curves were obtained by interpolating each fold’s curve onto 501 equally spaced false-positive rates; bands show the across-fold standard deviation. Fold distributions and bands describe repeated-validation variability, rather than confidence intervals from independent samples. For clade/PCAcluster-specific performance, metrics were calculated separately for each cluster using its held-out predictions in each repeat of grouped cross-validation, then averaged across the five repeats. Balanced accuracy was undefined for clusters containing only one phenotype class; recall was undefined without resistant isolates and precision without predicted-resistant isolates.

### Within-group random-forest feature importance

To identify predictive genes within genetic backgrounds, gene-level variant random forests were fitted separately in clade 1 (n=271; 167 resistant), clade 3 (n=63; 21 resistant), clade 4 (n=62; 10 resistant) and PCA cluster 6 (n=63; 12 resistant). Each group used the corresponding rows of the gene-level input matrix described above; variant eligibility and the initial gene-frequency filter were not recalculated within groups. Feature screening, forest settings, three-fold inner hyperparameter tuning and threshold selection followed the common prediction procedure. Outer validation comprised five repeats of five-fold stratified cross-validation for clade 1 and five repeats of three-fold stratified cross-validation for the other groups, producing 25 and 15 fitted outer-training models, respectively.

Gene importance was measured using the random forest’s mean decrease in Gini impurity^139^. Gini impurity describes how mixed the phenotype classes are within a tree node: G = 1 − pR² − pS², where pR and pS are the class-weighted proportions of resistant and susceptible training observations reaching that node. Impurity is zero when all observations belong to one class and greatest when the two classes are equally represented. A split is informative when it creates child nodes with lower weighted impurity than the parent. For each tree, the decreases attributable to splits on a gene were weighted by the fraction of training weight reaching each node, summed and normalized across features. The forest importance was the average of these normalized tree importances, normalized to sum to one across the fitted forest’s features.

For each genetic group, normalized gene importance was averaged across all outer-training models. A gene received zero for any model in which it was not retained, ensuring that the mean reflected both its contribution when selected and its consistency across folds. Genes were ranked by decreasing mean importance and plotted at their genomic midpoints. Gini importance is a relative measure of a feature’s contribution to the fitted classifiers: it has no direction of effect, is influenced by correlated predictors, and does not establish that the carried variants cause resistance.

The 20 highest-ranked genes from each group were used to construct a gene network using the annotation and network procedure described above, with genetic groups replacing antifungal drugs. Connections were coloured by group, and the gene-marker area reflected the number of group-specific top-20 lists containing that gene. Functional descriptions were obtained and reviewed using the same Candida Genome Database and primary-literature sources. For this fluconazole-focused network, magenta denoted published azole-response or target evidence, whereas orange denoted evidence involving other drugs, cell-wall biology or explicitly qualified similarity hypotheses. Black and grey retained the previously defined meanings. The same evidence categories identified labelled top-20 genes in the importance plots.

### Machine learning with variant effect predictors (VEPs) and FungAMR

For the machine learning with VEPs and FungAMR, only samples with a substitution in at least one of the proteins reported in FungAMR and from PCA clusters with n>15 were included (seven clusters totaling R=163 and S=219). Samples with the same genotypes were also removed to avoid leakage.

FungAMR confidence scores were rescaled to obtain a linear scale improving the ability of the random forest to distinguish between resistance and susceptibility. Positive scores correspond to evidence for resistance, with a score of 8 representing the highest confidence (originally 1 in the database), and a score of 1 representing the lowest confidence (originally 8). Similarly, negative scores correspond to evidence for susceptibility, with -8 indicating the highest confidence (originally -1) and -1 the lowest (originally -8). For mutations with both resistance and susceptibility scores for a given antifungal, only the score with the highest confidence level was kept. Each strain was assigned only one confidence score since a single resistance mutation is enough to confer antifungal resistance. Accordingly, when a strain harbored multiple mutations, it was assigned only the highest resistance score. When no scores were available for a sample, the value of 0 was given.

Predictions of variant effects were computed for proteins reported in the FungAMR database that have previously been associated with antifungal resistance. Three variant effect predictors (VEPs) were used: FoldX^78^, GEMME^76^ and ESM^77^. The proteins included are Ben1, Cap1, Cdr1, Cdr2, Cmp1, Cnb1, Crz1, Erg1, Erg2, Erg3, Erg11, Fks, Fur1, Mkc1, Mkk2, Mrr1, Mrr2, Rlm1, Rnr1, Rta2, Swi4, Tac1 and Upc2.

The effects of amino acid substitutions on protein stability were estimated using FoldX with the available 3D structures from the alphafold database. The RepairPDB function was run ten times to repair incorrect torsion angles, VanderWaals clashes and total energy of the structures. We used the MutateX Python package to perform an in silico deep mutational scan of the entire protein. The BuildModel function from FoldX was applied to compute the change in Gibbs free energy (ΔΔG) between each mutant and the wild type.

We used GEMME and ESM models to predict the functional impact of mutations. To run GEMME, we first produced a multiple sequence alignment with the orthologous sequences of interest as a query using HHblits function from HH-suite3. Initial HHblits iterative sequence search filter criteria were a minimum coverage of 80% with query sequence, at least 20% identity to the query and maximum 98% and the e-value cutoff for inclusion was 0.0001. Filter criteria were then gradually relaxed to get a minimum of 200 sequences in the final alignment. GEMME was then run locally through docker desktop to get the predicted effect of every single amino acid substitution across the proteins. We computed ESM scores using the ESM variant as described in Brandes et al. 2023, which is based on the ESM-1b protein language model. For each missense mutation, the variant effect score was computed as the difference in log-likelihood between the mutant and the wild-type amino acid at the same position.

For GEMME and ESM, lower scores indicate a stronger predicted impact, with 0 representing no predicted effect. For FoldX, positive ΔΔG values predict that a substitution will be destabilizing, with 0 representing no predicted effect. For each sample, the minimal GEMME and ESM scores and the maximal FoldX ΔΔG for each gene were used as features. Therefore, when multiple substitutions occur in the same protein, the score corresponding to the substitution with the largest predicted impact is used. If a sample carries no substitutions in a protein, a value of 0 was assigned for all VEPs.

A random forest classifier was trained using repeated nested cross-validation with VEPs alone and combined with FungAMR confidence scores as features. In the outer loop, model generalization was evaluated using Stratified K-Fold and Group Stratified K-Fold (five folds for five repeats = 25 total folds). Hyperparameters were optimized independently within each outer training set using GridSearchCV with 3-fold Stratified K-Fold CV in the inner loop. The hyperparameter search space included the number of trees (n_estimators: 50, 100, or 200), maximum tree depth (max_depth: 5, 10, or unrestricted), minimum samples required for a split (min_samples_split: 3 or 5), minimum samples per leaf (min_samples_leaf: 3 or 5), number of features considered at each split (max_features: √p, 50% of features or all features) and class weighting (class_weight: none or balanced). Model performance was assessed on the 25 outer test folds using balanced accuracy, precision, and recall. Feature contributions were assessed using SHapley Additive exPlanations (SHAP) with TreeExplainer on the test set for each cross-validation fold. SHAP values were extracted for the positive class (resistant). SHAP values were calculated independently for each outer cross-validation fold and then averaged across repeated evaluations of the same sample. The resulting aggregated SHAP values were used to visualize the global contribution of features to model predictions.

### Prioritizing candidate genes

We followed up genes from the GWAS and random-forest networks to identify associations with the R phenotype that persisted after accounting for genetic population structure and correlated carriage of other candidate genes. Analyses used drug-specific cohorts with a resolved absence classification for FungAMR resistance-associated substitutions carrying drug-matched positive resistance-confidence scores of 1–4. Isolates with unknown FungAMR status were excluded. Existing R phenotype and S phenotype classifications were retained. The resulting dataset contained 1,467 isolate–antifungal observations from 471 distinct isolates. Eligible R phenotype/S phenotype sample sizes were 62/268 for fluconazole, 45/108 for voriconazole, 63/79 for posaconazole, 9/374 for caspofungin, 1/192 for anidulafungin and 2/264 for micafungin. An isolate could contribute to multiple antifungal cohorts.

The candidate set comprised 91 GWAS-network genes evaluated across six antifungals and 76 random-forest-network genes evaluated across the three azoles. Overlapping gene–antifungal hypotheses were counted once, yielding 759 hypotheses across 162 distinct genes. Variant eligibility was inherited from the original drug-specific gene-carriage definitions: predicted HIGH- or MODERATE-impact alleles, including qualifying gene-disrupting annotations, with carrier frequency >0 and ≤5% in the original cohort. Eligibility was not recalculated after FungAMR filtering. Gene carriage was encoded as the presence of at least one eligible alternate allele, allowing different rare variants within a gene to contribute to the same gene-level indicator. A positive call took precedence over missing calls. If no eligible variant was detected but at least one eligible call was missing, gene-carriage status was classified as unknown; non-carriage required all eligible calls to be known and absent. Unknown calls were excluded separately for each analysis and were never imputed as absence. No additional minimum number of carriers or called isolates was imposed.

Initial gene enrichment was assessed using two-sided exact conditional tests stratified by the original principal component analysis (PCA) clusters. Stratification accounted for genetic population structure, while exact inference avoided reliance on large-sample approximations for sparse carriage counts. Tests estimated a common odds ratio comparing gene carriage between the R phenotype and S phenotype classes. Fisher’s exact test was used when only one informative stratum remained. A stratum was informative when it contained both phenotype classes and both carriage classes. Benjamini–Hochberg correction was applied across the fixed family of 759 hypotheses, and genes qualified for follow-up when the odds ratio exceeded 1 and the adjusted P value, q, was <0.05. Study was not included as an adjustment variable.

To assess whether initial associations could reflect correlated carriage, candidate partners for each target gene were restricted to other qualifying genes for the same antifungal. For each gene pair, we used isolates with known calls for both genes, pooling the two phenotype classes. Each binary carriage variable was centred by subtracting its mean within each PCA cluster, calculated on that pair’s jointly called subset. Pearson correlation was then calculated across the pooled residuals. This approach selected partners according to carriage relationships remaining after removal of differences in average carriage between genetic clusters. Partners with absolute correlation ≥0.20 were included, allowing both positive and negative correlations. This operational threshold was assessed in sensitivity analyses using thresholds of 0.15, 0.25 and 0.30.

For each target gene, the joint analysis retained only isolates with known calls for the target and every selected partner. Exact conditional tests stratified simultaneously by PCA cluster and the complete binary carriage pattern of all selected partners, thereby accounting for those partners jointly. A PCA-stratified comparison using the identical isolate subset was also calculated to distinguish changes associated with conditioning from those associated with sample exclusion. Follow-up q values were calculated across the fixed family of 36 target-gene hypotheses, separately for each correlation threshold and adjustment scheme. Genes were prioritized for individual-variant follow-up when they had at least one selected partner, a conditional odds ratio >1 and q <0.05. Associations without selected partners were not treated as evidence of successful co-carriage adjustment. Tests without informative strata were classified as non-estimable. Across all correction families, non-estimable tests contributed P = 1 solely for multiple-testing correction, while their reported P and q values remained missing. Boundary odds-ratio estimates of zero or infinity were retained when exact inference was available.

Individual-variant analyses evaluated 1,692 eligible variant–antifungal records across the initially qualifying gene–antifungal families. Each variant was tested using its own subset of isolates with known calls, with two-sided exact inference stratified by PCA cluster. Benjamini–Hochberg correction was performed separately within each gene–antifungal family. All variants with odds ratio >1 and within-family q <0.05 were displayed for genes meeting the joint co-carriage prioritization criteria; no limit on the number of displayed variants was applied. Individual-variant tests did not adjust for co-carried genes: co-carriage adjustment applied to gene selection. Carrier percentages used the number of isolates carrying the variant divided by the number with a known call, separately for each phenotype, with 95% Wilson binomial confidence intervals.

The isolate-level carriage display included R phenotype isolates only and was descriptive; enrichment tests compared both phenotype classes, and partner selection pooled both classes. Gene and partner selection used the same discovery data, so these analyses were exploratory follow-up rather than independent validation. Conditional associations describe evidence remaining after adjustment for the selected partners and do not establish independence from every other genetic factor.

